# Subcellular transcriptome sequencing with single cell APEX-seq identifies regulators of cell-cell interactions

**DOI:** 10.64898/2026.03.17.712496

**Authors:** Andrew Xue, Bo Cai, Qian Xue, Nianping Liu, Xiaojie Qiu, Rogelio A. Hernández-López, Alice Y. Ting

**Affiliations:** Department of Genetics, Stanford University, Stanford, CA, 94305, USA; Biophysics Program, Stanford University, Stanford, CA, 94305, USA; Department of Bioengineering, Stanford University, Stanford, CA, 94305, USA; Department of Computer Science, Stanford University, Stanford, CA, 94305, USA; Departments of Biology, and by courtesy, Chemistry, Stanford University, Stanford, CA, 94305, USA; Biohub, San Francisco, CA, 94158, USA; Parker Institute for Cancer Immunotherapy, San Francisco, CA, 94129, USA

## Abstract

Single-cell RNA sequencing has transformed our understanding of tissue complexity and heterogeneous cell states, yet provides little information about the subcellular organization of transcriptomes - despite the central role of RNA localization in splicing, translation, and function. Here we introduce single-cell APEX-seq (scAPEX-seq), a proximity labeling-based method for mapping subcellular transcriptomes at single-cell resolution. Improvements in probe design and RNA recovery enable APEX integration with droplet-based RNA-seq to capture endoplasmic reticulum–associated transcripts from thousands of individual cells. Applied to tumor–macrophage co-cultures, ER-targeted scAPEX-seq revealed interaction-dependent cell states and transcriptomic signatures by enriching for cell surface and secretory transcripts that are poorly resolved by conventional scRNA-seq. We further applied scAPEX-seq to short- and long-term co-cultures of HER2+ tumor cells with human chimeric antigen receptor (CAR) T cells, resolving distinct activated CAR T cell states, including populations characterized by upregulated NT5E or CTSW expression. We showed that overexpression of CTSW, a cathepsin protease, in CAR T cells promotes stem-like phenotypes, long-term proliferation, and sustained tumor cell killing. scAPEX-seq provides a powerful and scalable approach for profiling subcellular RNA populations, enabling the discovery of cell-cell interaction regulators missed by conventional approaches.

## Introduction

The RNA content, or transcriptome, of living cells is a signature of both cell identity and cell state. Technologies for transcriptome measurement are therefore highly valuable for understanding the biology of cells and organisms. Over the past decade, two parallel advances have dramatically improved the power of RNA sequencing: single-cell transcriptomics and subcellular transcriptomics.

The explosion of single cell technologies has led to a deeper understanding of tissue architecture, organ development, tumor microenvironments, and even how cell types evolve. Cell states that were not visible in averaged, bulk RNA-seq data can now be detected, classified, and studied. Furthermore, the proximity of cell types to each other, as revealed by spatial transcriptomic methods, sheds light on possible cell-cell interactions (CCIs).

Subcellular RNA analysis is important because RNA location is just as relevant as abundance, determining when and how an RNA is spliced, post-transcriptionally modified, translated, and degraded. As an example, NET1 mRNA localized to the cell periphery is translated into protein product that binds to membrane scaffolding proteins and functions in cell migration. However, when NET1 mRNA is localized to the nuclear periphery, its protein product binds to importin and functions in the nucleus^1^. Many RNA subpopulations are targeted to distinct organelles for local translation by ribosomes stationed there^2-4^. RNA mislocalization is increasingly recognized as a hallmark of disease^5^, such as in spinal muscular atrophy where the failure of β-actin mRNA to reach axonal processes leads to neurodegeneration^6^.

Therefore the ability to combine these two advances—subcellular transcriptome analysis at single cell resolution—could be transformative for peering deeper into cell states and understanding how RNA organization controls cell form and function. For example, the selective analysis of transcripts localized to the ER membrane that encode cell surface and secreted proteins could illuminate mechanisms of cell-cell interactions, and how these change under different conditions and in varying environments.

To date, the best methods for such analysis, such as MERFISH^7^, seqFISH^8^, and STARmap^9^, are based on probe hybridization and microscopy. Though powerful, these imaging-based methods are limited by their spatial resolution (typically ~1 µm), sensitivity, and throughput; they also require a pre-selected set of transcripts that are targeted by complex probe designs. It is not currently possible, for example, to use these methods for unbiased profiling of the ER-associated secretory transcriptomes of single cells.

To overcome these limitations, an unbiased, sequencing-based rather than hybridization-based approach to subcellular RNA analysis with single cell sensitivity is needed. The most widely-used method for bulk subcellular RNA analysis – biochemical fractionation of organelles followed by RNA-seq – cannot be performed on single cells. As an alternative, we explored the use of APEX-seq^10^, a proximity labeling method for mapping subcellular transcriptomes in living cells. In this approach, the promiscuous peroxidase APEX2 is genetically targeted to subcellular compartments of interest, where it catalyzes 1-minute biotinylation of endogenous transcripts within ~10 nm for subsequent identification by RNA sequencing. Using bulk APEX-seq, we created an averaged spatial map of the human transcriptome in HEK 293T cells^10^. To marry APEX-seq with the benefits of single cell analysis, however, we needed to overcome two major hurdles: low RNA capture efficiency, and the challenge of enriching biotinylated over non-biotinylated transcripts at the single cell level.

In this work, we show that an overhaul of APEX-seq and careful integration with the 10x Genomics droplet microfluidics workflow enables sensitive and compartment-specific mapping of subcellular transcriptomes in single mammalian cells. By using single cell APEX-seq (scAPEX-seq) at the ER membrane of cocultured tumor cells, macrophages, and Chimeric Antigen Receptor (CAR)-expressing T cells, we discover subcellular transcriptomic profiles that reflect cell-cell interactions missed by conventional scRNA-seq. Our work leads to the discovery that Cathepsin W overexpression drives CAR T cell proliferation and persistence, improving its capacity to kill tumor cells over repeated cycles of exposure. Overall, scAPEX-seq offers a versatile strategy for mapping the subcellular transcriptomes of multiple cell types and for uncovering intercellular interactions missed by conventional approaches.

## Results

### Development of APEX-seq2

First-generation APEX-seq^10^ used biotin-phenol (BP) to label RNAs proximal to APEX before cell lysis and streptavidin enrichment of biotinylated transcripts. However, APEX-seq is limited by its low recovery of compartment-specific RNAs (~0.1%), necessitating a large number of input cells (~10^6^ cells) to obtain sufficient material for sequencing. We aimed to improve the sensitivity of APEX-seq in order to lay the groundwork for single cell analysis.

We first simplified the APEX-seq workflow by omitting steps for RNA elution from streptavidin beads and subsequent RNA purification. Instead, we amplified RNAs directly on the beads (**Figs. S1A, S1B**), which improved the recovery of biotinylated transcripts by 4-7 fold when using HEK 293T cells expressing APEX targeted to the mitochondrial matrix (**Fig. 1C**). Next, we considered the BP probe, which must be pre-incubated with cells for 30 minutes at high concentration (500 µM) due to its poor membrane permeability. In certain cell types, such as Raw 264.7 macrophages, BP entry is so poor that no enrichment of target RNAs is observed (**Fig. S1C**). Recently, we reported that the alternative APEX substrate alkyne-phenol^11^ gives superior protein labeling and enrichment due to its improved entry into cells. Alkyne probes, however, require copper-mediated Click chemistry for derivatization which can be damaging to RNA^12^. To circumvent this, we synthesized and tested a phenol-azide probe (PA) to replace BP (**Fig. 1B**). PA, in contrast to alkyne-phenol, can be derivatized under mild, copper-free conditions using strained alkynes such as dibenzocyclooctyne (DBCO)^13^. In a side-by-side comparison, replacing 500 µM BP with just 50 µM PA followed by DBCO Click improved RNA recovery by 2-3-fold (**Fig. 1C**).

**Figure 1.**
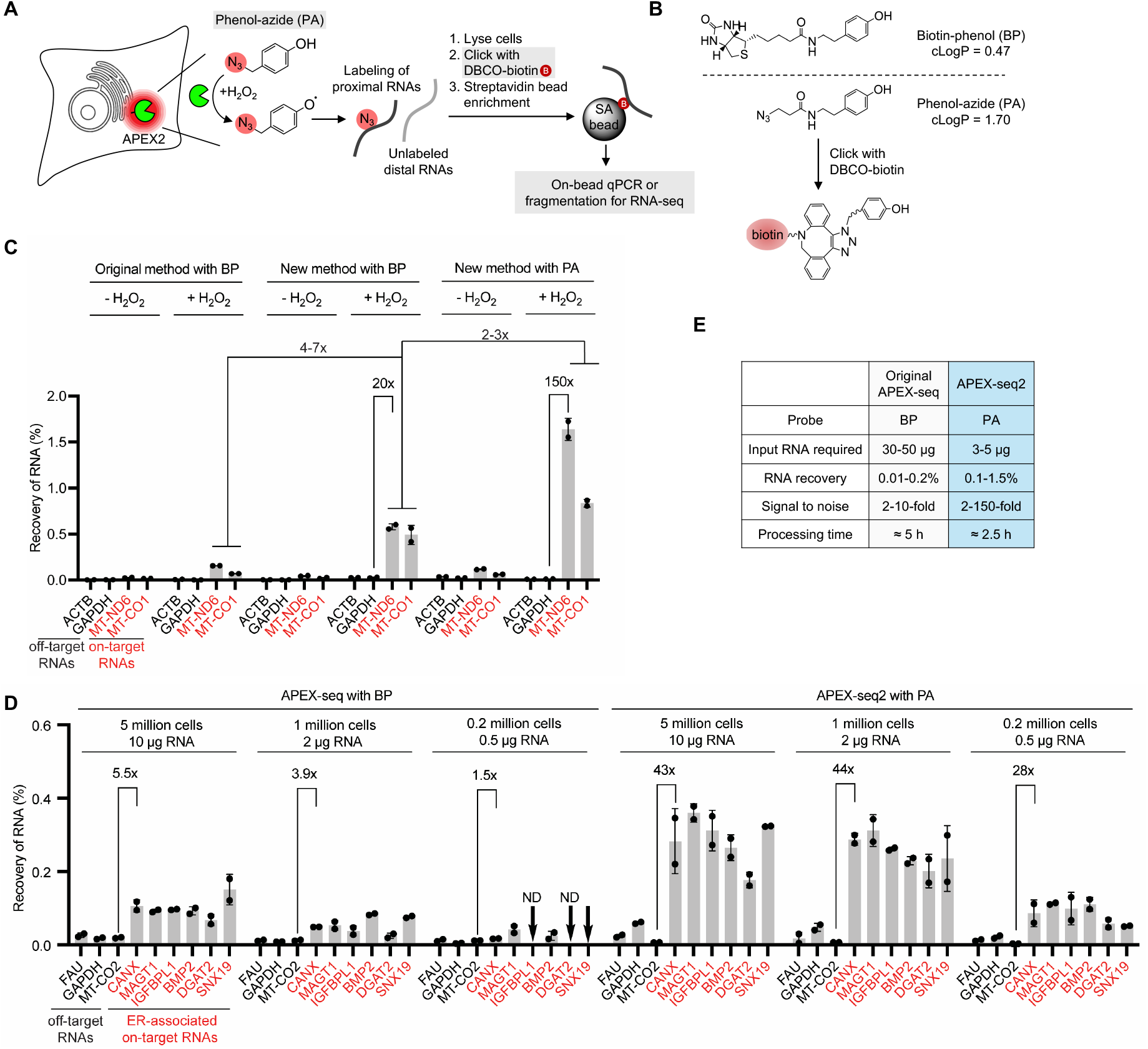
Improved transcript recovery with APEX-seq2. (**A**) Schematic of APEX-seq2 protocol. APEX2 (green) is genetically targeted to a subcellular region of interest, such as the ER membrane facing cytosol. Incubation with phenol-azide and H_2_O_2_ results in labeling of proximal endogenous transcripts with an azide functional group (red). After cell lysis, labeled transcripts are derivatized with DBCO-biotin and enriched using streptavidin beads. Differences from original APEX-seq protocol^10^ are shaded grey. (**B**) Structures of biotin-phenol (BP) and phenol-azide (PA). PA is derivatized with DBCO-biotin via strain-promoted azide-alkyne cycloaddition, enabling copper-free streptavidin enrichment of tagged RNAs. (**C**) RNA recovery using original APEX-seq vs. APEX-seq2. HEK293T cells expressing APEX2 targeted to the mitochondrial matrix were labeled with BP or PA for 1 minute. On-target mitochondrial (red) and off-target non-mitochondrial (black) transcripts were quantified by qRT-PCR before and after streptavidin enrichment, to determine fraction of RNA recovered. n = 3 experiments, 2 replicates per condition. Data are mean ± s.d. (**D**) RNA recovery with different cell and RNA inputs. Labeling was performed on 0.2 to 5 million HEK293T cells expressing APEX2 targeted to the ER membrane. Following RNA isolation, 10 ug, 2 ug, or 500 ng RNA were used for streptavidin enrichment. Transcript recovery was calculated as in (C). n = 2 experiments, 2 replicates per condition. Data are mean ± s.d. ND, not detected. (**E**) Comparison of original APEX-seq^10^ and APEX-seq2. More details in **Fig. S1.**

Combining these two modifications in “APEX-seq2” (**Fig. 1A**), we assessed the enrichment of mitochondrial transcripts by APEX2 targeted to the mitochondrial matrix of HEK 293T cells. **Figure 1C** shows that RNA recovery was ~1%, 10-fold higher than that obtained using the original method with BP. Furthermore, specificity was excellent, with on-target mitochondrial transcripts enriched by >100-fold over off-target cytosolic markers. To evaluate the sensitivity, we performed RNA enrichment using progressively fewer input cells. 200K to 5 million HEK 293T cells expressing APEX2 at the ER membrane were labeled with PA for 1 minute and lysed, yielding 500 ng to 10 μg of total RNA, respectively. After copper-free Click with DBCO-biotin and streptavidin enrichment of biotinylated transcripts, we performed RT-qPCR analysis. APEX-seq2 gave 28-fold enrichment of ER-associated secretory RNAs over cytosolic off-target RNAs, even with the lowest RNA input (500 ng). The original method was unable to enrich any secretory RNAs on this scale (**Fig. 1D**). Overall, our protocol modifications improved both RNA recovery and sensitivity of APEX-seq without a loss of specificity (**Fig. 1E**). In addition, the total process time of APEX-seq2 was reduced by half (**Fig. S1B**).

### Improved mapping of subcellular transcriptomes using bulk APEX-seq2

To test APEX-seq2 more comprehensively, we generated HEK293T cells stably expressing APEX2 targeted to five distinct subcellular compartments: the cytosol, ER membrane (ERM) facing cytosol, outer mitochondrial membrane (OMM) facing cytosol, nucleus, and mitochondrial matrix. The localization and activity of each construct was confirmed by microscopy, using both PA and BP as probes (**Fig. 2A**). When we enriched PA-labeled transcripts for RT-qPCR analysis, we observed 10-100-fold more on-target RNAs than off-target RNAs in membrane-enclosed compartments such as the mitochondrial matrix and nucleus (**Fig. 2B**). In non-enclosed regions of the cell, such as the ERM or OMM facing cytosol, on-target transcripts were still enriched, though to a lesser extent than closed-compartment transcripts (**Fig. 2B**).

**Figure 2.**
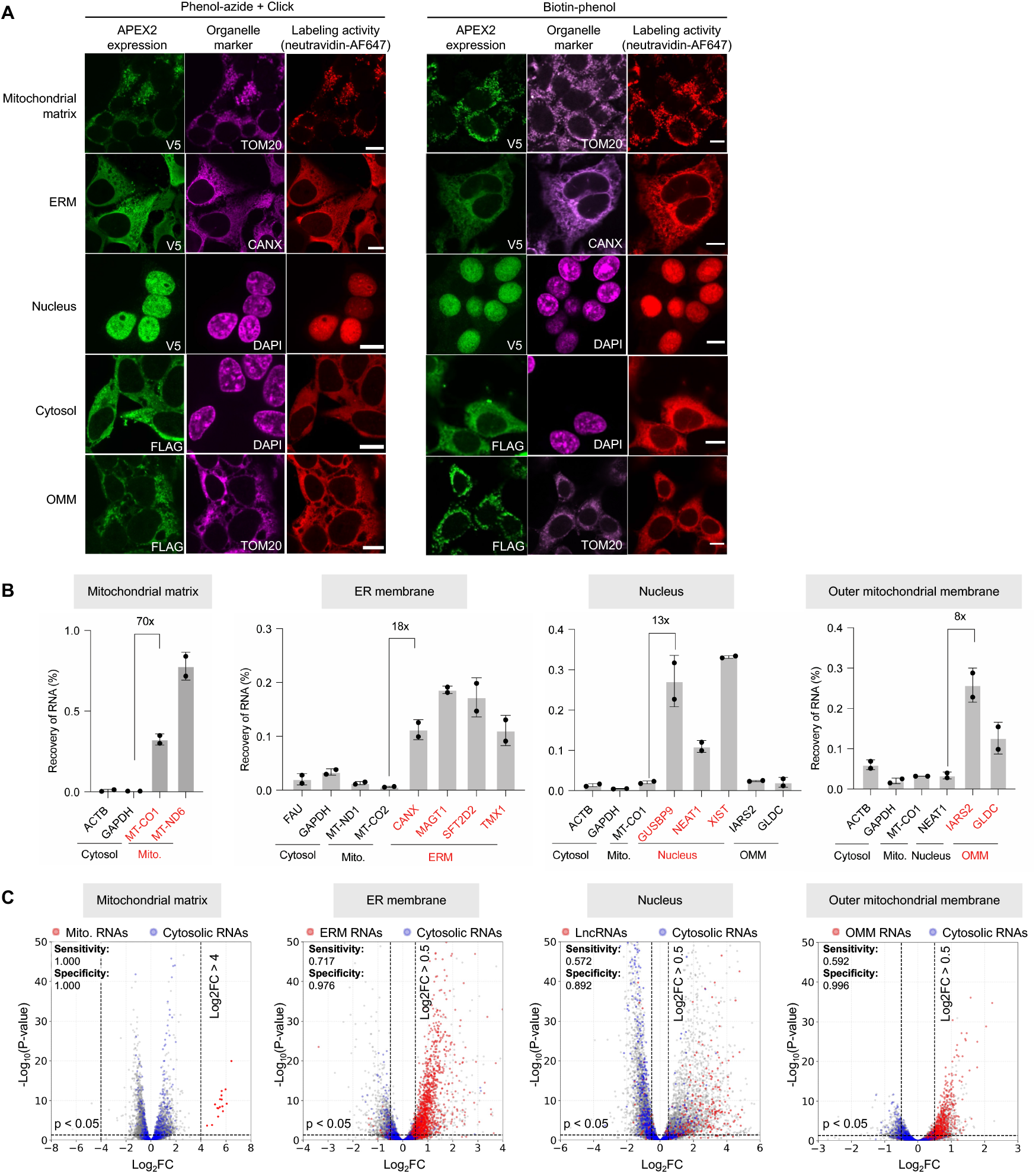
Enrichment of organelle-specific transcriptomes by bulk APEX-seq2. (**A**) Confocal fluorescence imaging of APEX2 localization and labeling activity in various subcellular compartments. HEK293T cells stably expressed the indicated APEX2 construct were labeled live with PA (left) or BP (right) for 1 minute, then fixed and Clicked with biotin-alkyne (for PA). Cells were stained with neutravidin-AF647 (red) to detect biotinylation, anti-V5 or anti-FLAG antibodies to detect APEX expression (green), and an antibody against an organelle marker (purple). Scale bars, 10 μm. (**B**) Compartment-specific enrichment of RNAs by organelle-targeted APEX2 constructs. Cells from (A) were labeled and processed as in **Fig. 1A**. RNA recovery calculated from qRT-PCR before vs. after streptavidin enrichment. n = 2 experiments, 2 replicates per condition. Data are mean ± s.d. (**C**) Bulk APEX-seq2 sequencing in multiple compartments of HEK293T cells. Cells from (A) were labeled and processed as depicted in **Fig. 1A**. Detected transcripts are plotted by log2 fold-enrichment (L2FC) in the specified compartment versus cytosol. On-target RNAs are colored red and off-target RNAs are colored blue. Specificity is calculated as 1 - true negatives / (true negatives + false positives). Sensitivity is calculated as true positives / (true positives + false negatives). LncRNAs, long non-coding RNAs.

We then performed bulk sequencing of APEX-seq2 samples (**Fig. 2C**). All conditions used 10-fold less input material than in our original APEX-seq study^10^ (2 million cells, giving 4 μg total RNA). Differential enrichment analysis was performed using cytosolic APEX2 read counts as a control. In the mitochondrial matrix, APEX-seq2 enriched all 13 mRNAs and 2 rRNAs encoded in the mitochondrial genome by >20-fold. The spatial specificity in other compartments, including open ones, was also high, at 98% for ERM and 99% for OMM. Sensitivity, calculated as the percent recovery of established true positives, was lower (57% for the nucleus, and 72% for the ERM), but still slightly improved compared to our previous APEX-seq method (52% and 70%, respectively, when analyzed in the same manner). We conclude that the improved RNA recovery of APEX-seq2 allows transcriptome analysis with 1/10^th^ the input material, and no loss of spatial specificity or sensitivity. These improvements should be particularly beneficial for analyzing rare cell types or compartments with low RNA content, such as neuronal synapses.

### Development of single cell APEX-seq (scAPEX-seq)

To adapt APEX-seq2 for single cell analysis, we chose the 10x Genomics droplet-based scRNA-seq platform for integration with APEX because it is widely used and provides high sensitivity and throughput at a low cost per cell. The primary challenge was how to enrich biotinylated transcripts using streptavidin without scrambling the transcriptomes of individual cells together. To achieve this, we first barcoded single cell transcriptomes using Gel Beads-in-Emulsion (GEMs) via a single cycle of reverse transcription without PCR (first strand synthesis in **Fig. 3A**). This produces full-length RNA-cDNA hybrids that carry both a cell-specific barcode and the PA labeling status from the APEX tagging step, because no PCR amplification has occurred yet. Then, we broke open the emulsions and pooled RNA-cDNA hybrids from different cells together for DBCO-biotin Click and streptavidin bead enrichment. Finally, both the streptavidin-bound transcripts and the supernatant transcript population were separately PCR-amplified to produce libraries for next-generation sequencing.

**Figure 3.**
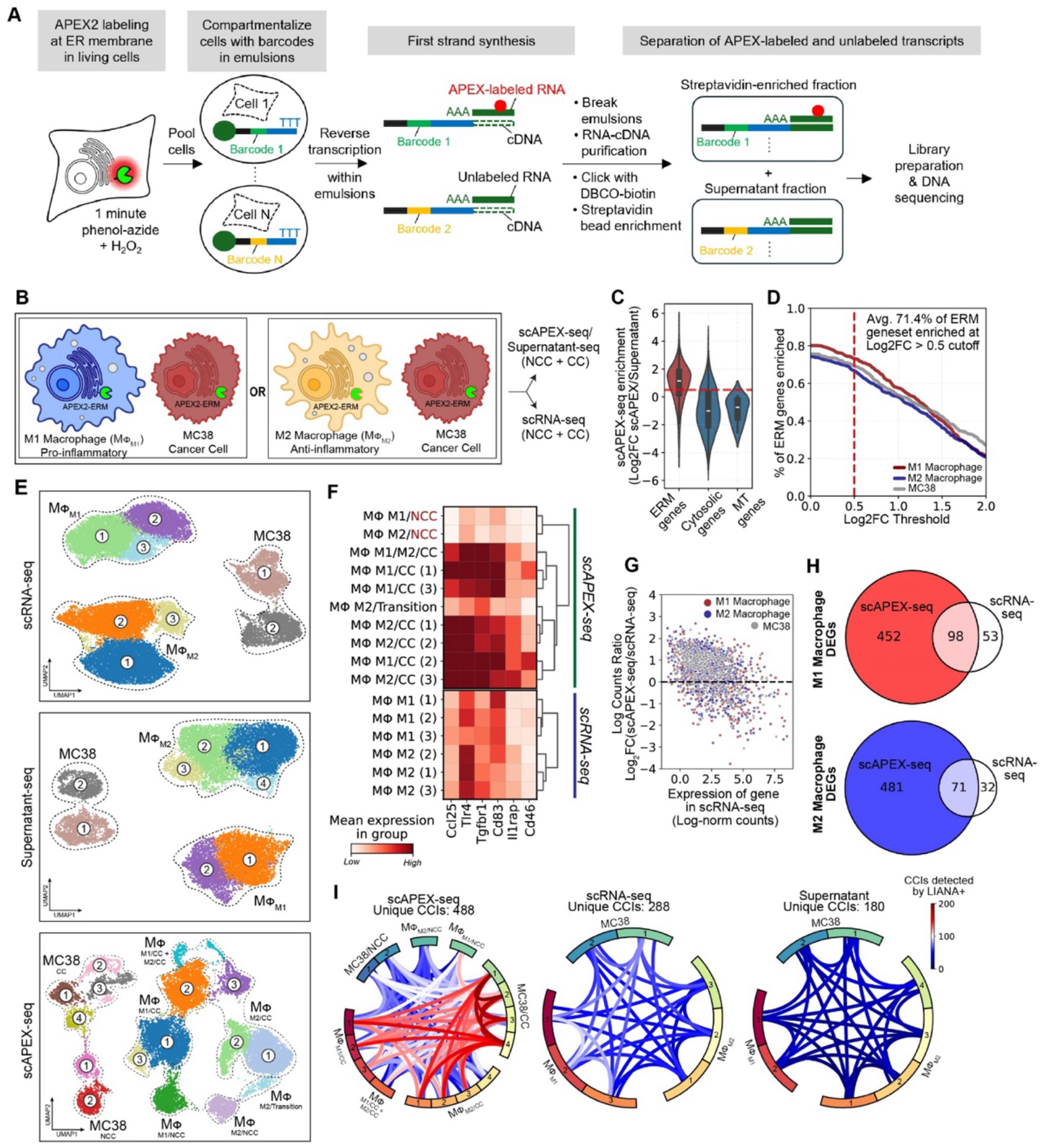
scAPEX-seq profiling of secretory transcriptomes in macrophage-cancer cell co-cultures. (**A**) scAPEX-seq workflow. After 1 minute of APEX2-catalyzed labeling with PA, cells are barcoded via first strand synthesis, then pooled for Click derivatization and streptavidin enrichment of biotinylated RNA-cDNA hybrids. Both the unlabeled supernatant fraction and the APEX2-labeled enriched fraction are amplified separately and processed for sequencing. (**B**) Co-cultures analyzed by scAPEX-seq and conventional scRNA-seq. RAW264.7 macrophages were polarized into either the M1 state or M2 state before co-culture (CC) with MC38 cancer cells for 16 hours. In non-coculture (NCC) control samples, cells were labeled separately with PA and H_2_O_2_, then quenched and combined before processing for scAPEX-seq as in (A). (**C**) Evaluation of scAPEX-seq spatial specificity. Previously reported ERM-localized RNAs, cytosolic RNAs, and mitochondrial RNAs are plotted by scAPEX-seq/Supernatant-seq counts. Red line denotes 0.5 Log2FC cutoff. (**D**) Evaluation of scAPEX-seq sensitivity. Fraction of 1593 true positive secretory mRNAs enriched at different L2FC cutoffs. (**E**) UMAP clustering of single cell data from samples in (B). Clusters are labeled by cell type, M1 versus M2 polarization, and co-culture versus non-coculture status based on established markers (see Methods). (**F**) Expression of coculture-specific markers across scAPEX-seq and scRNA-seq macrophage clusters. (**G**) scAPEX-seq sensitivity for low-count secretory genes. Log2 scAPEX-seq/scRNA-seq counts ratio plotted against gene abundance (in scRNA-seq) and colored by cell type. Line at Log2FC = 0 represents equal sensitivity. (**H**) Differentially-enriched genes (DEGs) discovered between different macrophage clusters discovered in each sample (scAPEX-seq vs. scRNA-seq). In scAPEX-seq, calculations were performed between CC and NCC clusters, while in scRNA-seq calculations were performed between the two largest clusters per cell type, as no clear NCC/CC clusters were resolved. (**I**) Chord diagrams of CCIs calculated by LIANA^29^ for scAPEX-seq, scRNA-seq, Supernatant-seq samples, depicting the number of unique and significant CCIs detected between clusters. Plots are shown on the same color scale.

Initially, we performed the DBCO-biotin Click reaction directly in broken emulsions, which contains a complex mixture of proteins, RNAs, and other cellular components. This resulted in low capture of unique molecular identifier (UMI) counts from the enriched fraction, even when using the improved PA-based labeling strategy. When we introduced a precipitation step to clean up the RNA-cDNA hybrids prior to the Click reaction (while preserving hybrid integrity), the UMI recovery increased by almost 100-fold (**Fig. S1D**). We also found that by performing reverse transcription before Click-derivatization of azide (which makes the moiety larger, impeding the reverse transcriptase) rather than post-Click as in bulk APEX-seq2, our average RNA recovery increased from ~1% to 1.5-3% (**Fig. S1E**).

For our first test of scAPEX-seq, we used co-cultures of tumor cells with macrophages (**Fig. 3B**). Macrophages are phagocytic immune cells that engage in complex bidirectional relationships with tumor cells^14^. Depending on context, they can either directly kill or initiate anti-tumor responses^15^, or they can enhance tumor progression and immunosuppression through conversion into tumor-associated macrophages (TAMs)^16^. Understanding the inter- and intracellular signaling pathways that control macrophage differentiation could inform the development of novel anti-cancer therapies.

Previous scRNA-seq studies of tumor-macrophage systems have successfully classified the major cell types and states^17,18^. However, inferring cell-cell interactions (CCIs) from this data is challenging, partly because transcripts encoding ligands, receptors, and co-receptors that mediate CCIs tend to be low in abundance^19-21^ (for example, the critical immune checkpoint marker PD-L1 is often missed^22,23^). In many cases, surface protein measurements like CITE-seq are multiplexed with sequencing to address this issue^24-26^. Another important limitation of scRNA-seq is that protein abundance, especially at the cell surface, is not well-correlated with total RNA levels^24,27^. Numerous mechanisms, including mRNA relocalization, translational regulation, post-translational modification, and protein secretion mechanisms play a role. These factors make the extraction of CCI information from conventional scRNA-seq data challenging.

We hypothesized that by applying scAPEX-seq with APEX targeted to the ER membrane, we might improve the detection of macrophage-tumor CCIs by focusing attention on the subset of genes most relevant to CCIs – those encoding cell surface and secreted proteins. The majority of such mRNAs are localized to the ER membrane, where they are translated by ER-associated ribosomes before translocation of their protein products across the ER membrane to the cell surface.

To perform the experiment, we first stably expressed APEX2-ERM in a murine colon adenocarcinoma cancer cell line (MC38) and RAW264.7 macrophages. Correct APEX2 localization and biotinylation activity were confirmed by confocal microscopy (**Fig. S2A**). We titrated PA probe concentration and analyzed enrichment of secretory vs. non-secretory transcripts by qRT-PCR to determine optimal labeling conditions (**Fig. S2B**). We also tested polarization of the macrophages into pro-inflammatory or anti-inflammatory states using LPS + IFN-γ or IL-4 + IL-13 treatments, respectively (**Fig. S2C**).

MC38 tumor cells were co-cultured (CC) with either pro-inflammatory or anti-inflammatory macrophages for 16 hours in total. In parallel, non-co-cultured (NCC) controls were prepared from tumor and macrophage cells cultured separately but mixed together immediately before 1-minute APEX labeling (**Fig. 3B**). CC and NCC samples were pooled and processed for scAPEX-seq. In addition, we collected the non-biotinylated supernatant fraction (“Supernatant-seq”) for sequencing. We also prepared barcoded whole-transcriptome samples to enable a direct comparison to conventional scRNA-seq.

### scAPEX-seq improves detection of cell-cell interaction-dependent cell states

We obtained high-quality single-cell transcriptome profiles for ~20k cells each from scAPEX-seq, Supernatant-seq, and scRNA-seq samples. The datasets showed a comparable number of features and UMIs per cell (**Fig. S3A**). We observed the three major cell types (tumor cells, pro-inflammatory macrophages, and anti-inflammatory macrophages), and enrichment of canonical markers in each group (**Fig. S3B**).

To assess the specificity of scAPEX-seq, we calculated the fold-enrichment of each gene in scAPEX-seq relative to Supernatant-seq using matched cell barcodes, as the datasets are derived from the same cell population. Secretory genes were strongly enriched relative to cytosolic and mitochondrial genes (**Fig. 3C**), suggesting that ER-targeted APEX labeling maintains spatial specificity at single cell resolution. To assess sensitivity, we quantified recovery of ER-annotated transcripts and found that scAPEX-seq enriched 70% of true-positive ER transcripts at a Log_2_FC of 0.5, similar to the sensitivity observed in bulk APEX-seq (**Fig. 3D**).

Supernatant-seq and scRNA-seq datasets were highly similar. Their UMAP projections showed relatively homogenous populations of each cell type, with similar subclusters (**Fig. 3E**). Integration of these clusters using Scanorama^28^ showed high overlap and correspondence across datasets (**Fig. S3C**). Gene expression signatures were also consistent between Supernatant-seq and scRNA-seq for tumor cells and for both macrophage types (**Fig. S3D**). Supernatant-seq likely provides a reasonable approximation of whole-transcriptome scRNA-seq because APEX labeling depletes a small fraction of compartment-specific transcripts from the supernatant.

The scAPEX-seq dataset was highly differentiated from both scRNA-seq and Supernatant-seq, with much greater cluster diversity and separation (**Fig. 3E**). Neither scRNA-seq nor Supernatant-seq could distinguish CC from NCC cells, whereas scAPEX-seq resolved these populations, based on established markers of coculture-induced expression (**Fig. 3F**). When all datasets were integrated using Scanorama^28^, Supernatant-seq and scRNA-seq clusters aligned closely as before (**Fig. S3E**). The NCC clusters from scAPEX-seq also integrated into the respective tumor cell, pro-inflammatory, and anti-inflammatory groups. However, CC clusters from scAPEX-seq remained distinct, indicating that scAPEX-seq captures interaction-dependent cell states that are obscured in whole-transcriptome measurements.

Two non-mutually exclusive mechanisms may underlie the enhanced sensitivity of scAPEX-seq to CCI-dependent states. First, scAPEX-seq recovers substantially higher counts for secretory transcripts than scRNA-seq, especially for low abundance genes (**Figs. 3G, S3F**). Second, scAPEX-seq detects CCI-induced changes in mRNA *localization* in addition to changes in transcript abundance. Using cell barcode matching, we calculated a localization ratio, defined as the ratio of scAPEX-seq counts to Supernatant-seq counts for each gene; this reflects the likelihood that a transcript is localized to the ER membrane (**Fig. S3G**). Many transcripts exhibited significant shifts in localization ratio between NCC and CC conditions, including numerous immunologically-relevant ones (e.g., TGFBR1 and CD83) (**Fig. S3H**). This observation highlights the importance of subcellular RNA organization in defining interaction-dependent cell states.

### scAPEX-seq identifies genes upregulated by tumor-macrophage interactions

We used our data to identify CCI-upregulated secretory genes that may play a role in driving or responding to tumor-macrophage intercellular signaling. Differentially-expressed genes (DEGs) between CC and NCC clusters were ~10-fold more numerous in the scAPEX-seq dataset than in the scRNA-seq dataset, with most of the latter being a subset of the former (**Figs. 3H, S3I-J**). We also applied LIANA+, which predicts ligand–receptor interactions from scRNA-seq data^29^, and found that scAPEX-seq produced many more unique interactions between cell types than scRNA-seq or Supernatant-seq did (**Fig. 3I**). The effect holds when using common cell-type labels alone (ignoring unique clusters that scAPEX-seq finds) (**Fig. S3M**). CCIs inferred by LIANA+ from scRNA-seq are largely a subset of those inferred from scAPEX-seq (**Fig. S3N**).

The interaction networks revealed by scAPEX-seq include canonical immunoregulatory and tumor–macrophage signaling pathways, such as Csf1-Csf1r, Jag1-Notch, and PD-L1-PD1 signaling (**Figs. S3K-L**). Multiple pathways were detected exclusively by scAPEX-seq, including the canonical Ccl2-Ccr2 and Ccl5-Ccr5 chemokine axes, TGFB1/2/3 signaling, Adam10/17 proteolytic interactions, damage-associated molecular pattern signaling (Hsp90b1-Tlr4/2), and the M2-promoting Grn-Tnfr2 axis. We also identified interaction pairs that have not previously been reported in the macrophage-tumor context, such as Fstl1-Dip2a^30^ (involved in anti-tumor immunity) and Pltp-Abca1^31^ (reported in macrophage STAT3/NFkB signaling). Consistent with these findings, scAPEX-seq revealed coculture-specific upregulation of Il34-Csf1r signaling (**Figs. S4A-B**), and corresponding downstream upregulation of Il10^32^ (**Fig. S4C**).

Overall, our findings from the application of scAPEX-seq to tumor-macrophage cocultures demonstrate that: (1) scAPEX-seq de-noises transcriptomic signals by enriching ER-associated secretory transcripts; (2) this improves the detection and analysis of CCI-dependent cell states and gene signatures; and (3) Supernatant-seq approximates the cell states revealed by whole-transcriptome scRNA-seq.

### scAPEX-seq applied to short-term (2 hour) tumor:CAR T-cell cocultures

Chimeric antigen receptor (CAR) T cell therapies have shown remarkable efficacy in hematologic malignancies, yet their performance in solid tumors remains variable and often limited by early dysfunction following tumor engagement. Even brief encounters with cancer cells can trigger incomplete activation, differentiation, and immunosuppressive programs that ultimately shape long-term CAR T cell persistence and efficacy. However, the early molecular events that occur immediately upon tumor contact, particularly those involving secretory and cell-surface signaling pathways, are poorly resolved by conventional scRNA-seq. We therefore applied scAPEX-seq to interrogate short-term tumor–CAR T-cell interactions, aiming to identify early regulators of CAR T-cell function.

We prepared 2-hour cocultures (CC) consisting of HER2+ breast cancer cells (HCC1569), human primary CD8^+^ anti-HER2 CAR T cells^33^, and non-CAR human primary CD8+ T cells, all expressing ERM-targeted APEX2 (**Figs. 4A** and **S5A-B**). CD8^+^ T cells lacking CAR served as an internal control to distinguish CAR-dependent from CAR-independent transcriptional changes. For this experiment, we leveraged a recent finding that metabolic incorporation of 6-thioguanosine (S^6^G) into RNA improves APEX2-mediated capture^34^ due to increased reactivity of the electron-rich thioamide moiety of S^6^G with electron-poor APEX-generated phenoxyl radicals (**Figs. S5C-D**). Cocultures were incubated with S^6^G for 2 h, labeled with PA for 1 min, quenched, and pooled for library generation. We also prepared a non-cocultured (NCC) control by culturing each cell type independently (also with S^6^G for 2 h) and pooling immediately before PA labeling.

**Figure 4.**
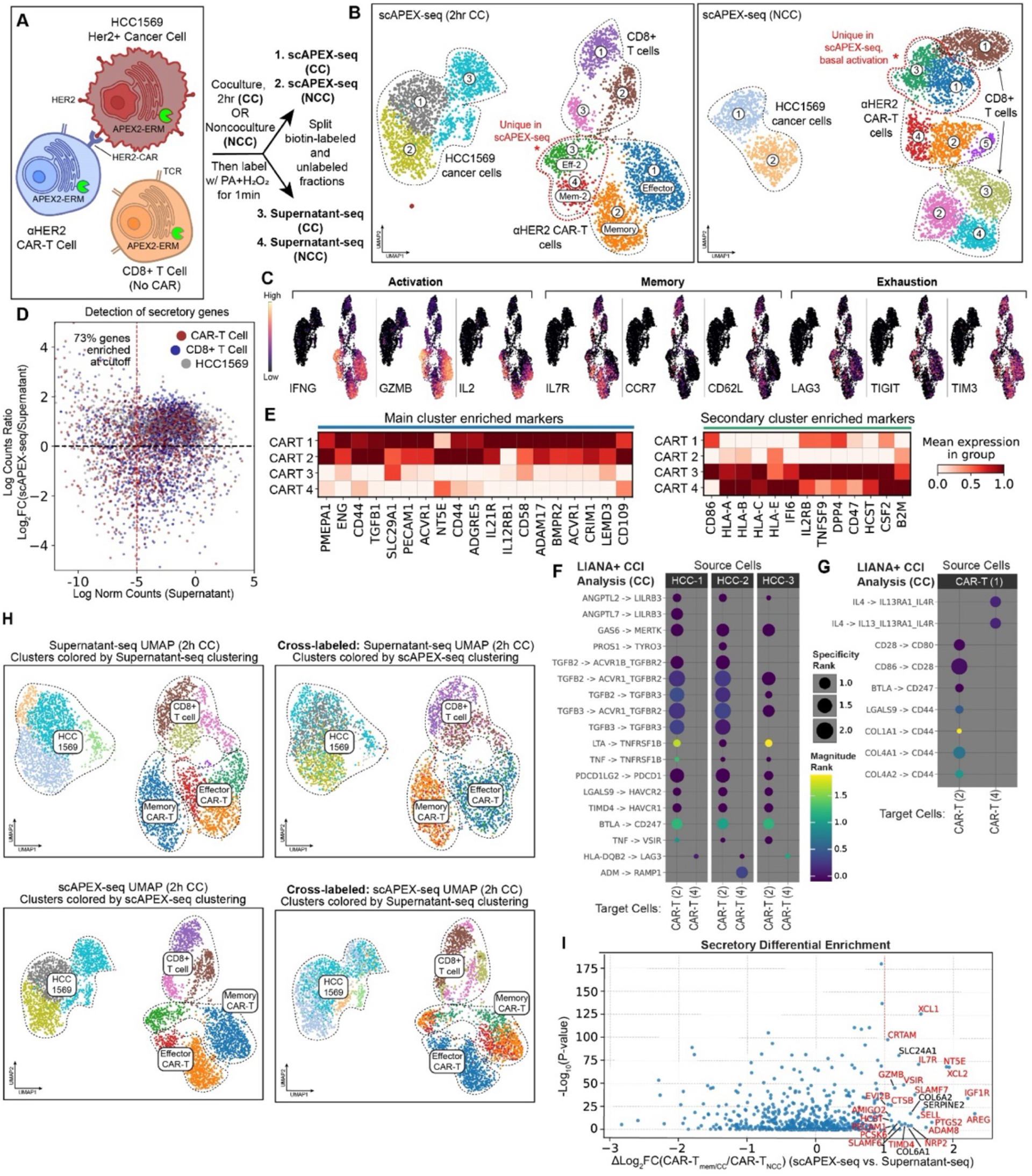
scAPEX-seq identifies secretory mRNAs involved in early CAR T cell activation. (**A**) Schematic of short-term CAR T cell-cancer cell coculture experiment. Three-cell cultures of HCC1569 cancer cells, untransduced CD8+ T cells, and anti-HER2 CAR T cells were incubated in a 6:1:2 ratio for 2 hours in the presence of 6-thioguanosine (S6G) to metabolically label newly synthesized RNAs. Cells were labeled with PA for 1 minute before processing for scAPEX-seq. (**B**) UMAP (Uniform Manifold Approximation and Projection) visualization of scAPEX-seq coculture (CC, left) and non-coculture (NCC, right) samples from (A). (**C**) Log-normalized expression levels of selected markers superimposed on scAPEX-seq UMAP. (**D**) Evaluation of scAPEX-seq sensitivity for low-count secretory genes. Log2 scAPEX-seq/Supernatant-seq counts ratio plotted against gene abundance (in Supernatant-seq) and colored by cell type. Line at Log2FC = 0 represents equal sensitivity. (**E**) Heatmap showing markers enriched in main vs. secondary CAR T clusters from scAPEX-seq. (**F**) Dot plot showing differential CCI detection by LIANA+ between cancer cells (source) and the main vs. secondary memory CAR T clusters (CAR T 2 and 4, respectively). Specificity and Magnitude ranks are LIANA+ derived aggregate metrics representing uniqueness and strength of a given interaction, respectively. (**G**) LIANA+ CCI detection between main effector CAR T cluster (source) and the main vs. secondary memory CAR T clusters (CAR T 2 and 4, respectively). (**H**) scAPEX-seq CC UMAP (bottom) and Supernatant-seq CC UMAP (top) with original labels compared to cells cross-labeled using matching barcodes to transfer cluster annotations between scAPEX-seq and Supernatant-seq samples. For example, for a given cell barcode A, it has both a cluster assignment based on scAPEX-seq clustering and Supernatant-seq clustering. Because cell barcode A is present in both samples, we can transfer the cluster assignment from scAPEX-seq and show it on the Supernatant-seq UMAP projection. (**I**) DEGs upregulated in memory CAR T cells compared to non-coculture CAR T cells in scAPEX-seq, normalized against the same comparison in Supernatant-seq. Genes previously implicated in T cell regulation and signaling are shown in red; genes without previous evidence are in black.

UMAP projections for the resulting datasets revealed distinct clustering of tumor cells, CAR T cells, and non-CAR T cells in both CC and NCC conditions (**Fig. 4B**). Strong enrichment of secretory transcripts was detected across all cell types when comparing gene counts in scAPEX-seq versus Supernatant-seq (**Fig. 4D**). In cocultured samples, CAR T cells, but not non-CAR T cells, exhibited robust induction of canonical activation markers including IFNG, GZMB, and IL2 (**Fig. 4C**).

The CAR T cells of the scAPEX-seq CC sample were grouped into 4 subclusters (**Fig. 4B**, left): a major effector and major memory cluster (1 and 2 in **Fig. 4C**), and secondary effector-like and memory-like clusters (3 and 4). The two major clusters exhibited hallmarks of classical CAR T cell activation, including upregulation of CD44/CD97, IL21R and IL12RB (downstream of 4-1BB signaling), and the costimulatory ligand CD58 (**Fig. 4E**). These populations also showed early signs of inhibition and immunosuppression, such as increased expression of ADAM17, a negative regulator of CD8^+^ T cell function, and multiple components of TGF-β signaling (TGFB1, BMPR2, ACVR1, CRIM1, LEMD3, CD109).

The secondary CAR T cell clusters displayed alternative activation states characterized by interferon signaling (HLA-A, HLA-B, HLA-C, HLA-E, B2M, and the interferon regulator IFI6), different immune activation markers (IL2RB, 41BBL, CD26), and survival-associated signals (HLA-E, CD47) (**Fig. 4E**). We hypothesized that these cells may represent a more interferon-sensitive population that promotes the activation of other T cells and is primed for survival.

LIANA+ analysis of CCIs involving these CAR T subclusters showed that immunosuppressive and checkpoint interactions, including TGFBRs, LILRBs, TIM-3, VISTA, and PD-1, were preferentially associated with the major memory cluster, but largely absent from the secondary memory cluster (**Fig. 4F**). Interactions between major effector and memory clusters reflected a balance of costimulatory and inhibitory signaling, whereas IL4 signaling was uniquely predicted in the secondary memory cluster (**Fig. 4G**) — consistent with partial Type 2 skewing^35^ and the observed upregulation of IL13 in this population.

Importantly, these patterns were not visible in the matched Supernatant-seq dataset. Instead of 4 distinct CAR T cell subpopulations, just one effector cluster and one memory cluster were observed (**Fig. 4H**). Transferring cluster labels between Supernatant-seq and scAPEX-seq, we found that scAPEX-seq labels were mixed within the Supernatant-seq UMAP projection, but not vice versa (**Fig. 4H**). This suggests that scAPEX-seq assigns a more refined set of cell-type labels and uncovers deeper structure than the Supernatant-seq sample. In the NCC scAPEX-seq samples, additional CAR T cell clusters were observed, with some showing signs of basal activation (upregulation of GZMB, IL2RA, and CD69) that may reflect tonic signaling, a known contributor to CAR T cell dysfunction (**Figs. S5E-F**).

Because memory states are associated with long-term persistence and favorable therapeutic outcomes^36^, we analyzed the memory CAR T clusters for DEGs. **Fig. 4I** shows transcripts that are enriched in scAPEX-seq relative to Supernatant-seq and in CC vs. NCC conditions. The majority of top-ranked genes have established roles in T cell differentiation and immune regulation. Among the top hits, we observed NT5E (CD73), an enzyme that catalyzes immunosuppressive adenosine production^37^, and SLC29A1 (ENT1), which is involved in adenosine transport into cells^38^. These findings point to early engagement of adenosine-mediated immunoregulatory pathways within memory CAR T cells. Follow-up experimentation on NT5E is described below.

### scAPEX-seq analysis of long-term CAR T cell response to repeat antigen stimulation

To model chronic tumor challenge, we established long-term cocultures in which anti-HER2 CAR T cells were repeatedly transferred onto fresh HER2-positive breast cancer cells (HCC1569) every 3 days for a total of 21 days (7 cycles). On day 21, both CAR T cells and surviving cancer cells were labeled with PA for 1 minute and processed for scAPEX-seq (**Figs. 5A, S5G**).

**Figure 5.**
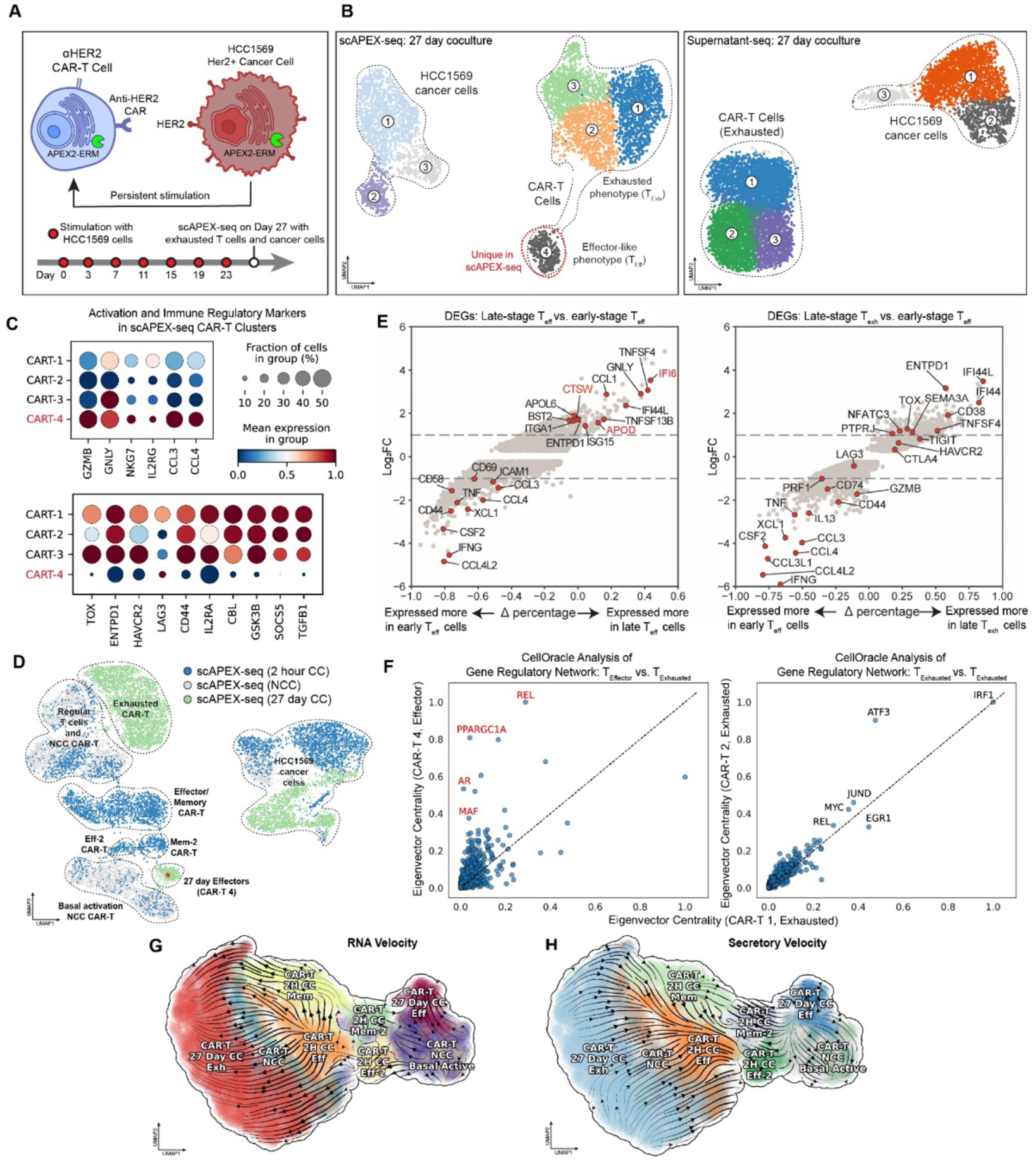
scAPEX-seq profiling of CAR T cell persistently stimulated with cancer. (**A**) Schematic of long-term CAR T cell-cancer cell coculture experiment. To anti-HER2 CD8+ CAR T cells expressing APEX2-ERM, HER2+ HCC1569 cancer cells were added repeatedly, every three days. On day 27, labeling was performed with PA for 1 minute, then cells were sorted (to remove debris) and processed for scAPEX-seq. (**B**) UMAP visualization of CAR T cells and cancer cells from experiment in (A). (**C**) Dot plots showing expression of selected activation-related and immune regulatory markers. Markers are on top are markers of activation that are upregulated in CAR T cluster 4 (effector-like) and downregulated in CAR T clusters 1-3 (with exhausted phenotype),while markers on the bottom represent markers downregulated in CAR T cluster 4 and include a panel of exhaustion markers. (**D**) Integration of long-term CAR T cell data with early CAR T cell activation data from **Fig. 4** using Scanorama^28^. Possible exhaustion-resistant cluster is at bottom (green, starred), next to early CAR T cell clusters in blue and grey. (**E**) Differentially expressed genes (DEGs) between late-stage T-effector cells and early-stage T-effector cells (left), and between late-stage T-exhausted cells and early-stage T-effector cells (right). CTSW in red was selected for validation in **Fig. 6**. (**F**) CellOracle^41^ analysis of gene regulatory networks in late T cell effector cluster versus exhausted CAR T cluster (left). On right is a control comparing two exhausted CAR T clusters. Previously identified exhaustion-related regulatory genes are colored red. (**G**) Dynamo^46^ modeling of transcriptomic vector fields in the early and late CAR T cell populations. Colors represent identified cell types. (**H**) Dynamo modeling^46^ of secretory vector fields using scAPEX-seq/Supernatant-seq enrichment ratios in place of RNA velocity in the early and late CAR T cell populations.

Supernatant-seq showed a relatively homogenous population of exhausted CAR T cells in the UMAP plot (**Fig. 5B**), but in scAPEX-seq we observed a unique and separate subpopulation of CAR T cells (**Fig. 5B**). This scAPEX-seq-specific cluster appeared to have comparatively higher effector function, as indicated by elevated levels of IFNG, GZMB, CCL3/CCL4, and NKG7 (**Fig. 5C**), and generally decreased expression of exhaustion markers (TOX, ENTPD1, HAVCR2, and LAG3)^39^. In addition, we observed decreased expression of CD25, TGFB1, and other immunoregulatory genes including SOCS5 and GSK3B, a kinase which has previously been linked to worsened CAR T cell exhaustion, persistence, and memory generation^40^. To contextualize this late-stage population, we integrated long-term scAPEX-seq data with short-term (2 hr) CC and NCC datasets. Remarkably, the scAPEX-seq-specific “late effector” CAR T-cell cluster aligned most closely with the secondary memory and effector CAR T cell clusters identified during early tumor interaction (**Fig. 5D**). In the integrated embedding, late exhausted CAR T cells align most closely with untransduced CD8 T-cells. We hypothesize that the “late T-cell effector” cluster may represent longer-surviving T-cells that retain the ability to kill tumor cells despite persistent antigen exposure.

To further characterize this putative persistent population, we performed DEG analysis, which showed upregulation of important cytotoxic and immune checkpoint genes compared to the exhausted late CAR T-cell cluster (**Fig. 5E**). We applied CellOracle^41^ to infer gene regulatory networks and identify transcription factors that may be controlling these cell states. Although APEX2-ERM preferentially captures secretory transcripts, counts are still detectable for a range of non-secretory transcripts including those encoding transcription factors. Comparing the network centrality of transcription factors in the late T cell effector cluster versus exhausted clusters, we observed significant shifts, which were not seen when comparing two different exhausted clusters (**Fig. 5F**). Several transcription factors predicted as highly important to our persistent cluster, such as AR, MAF, REL, and PPARGC1A, have previously been linked to regulation of T cell exhaustion, metabolism and persistence^42-45^.

To investigate the developmental origins of the late T cell effector population, we used Dynamo^46^ to infer cell state transitions across integrated early and late scAPEX-seq data using RNA velocity. RNA velocity trajectories suggest multiple distinct paths from early activation toward exhaustion or sustained effector function. We observe that early-stage effectors (CART-2H-EFF) are predicted to progress to exhaustion through a non-activated phenotype (CART-NCC), which may represent a gradual loss of CAR stimulation (**Fig. 5G**). CAR T cells that differentiate into a memory phenotype eventually progress towards exhaustion as well, but through a different pathway (**Fig. 5G**). The secondary memory-like cluster we identify by scAPEX-seq in the 2-hr sample presents an alternative fate, in which CAR T cells can differentiate towards the late effector subset of CAR T cells that we identified. Early NCC cells also have a path towards this late effector subset, possibly due to having low levels of activation/effector function without having accumulated exhaustion signals from antigen exposure. These results suggest that early divergence in CAR T cell state, which is detectable by scAPEX-seq but not Supernatant-seq, can predict long-term functional outcomes.

Finally, we asked whether changes in subcellular RNA localization might provide additional insight into CAR T cell state transitions. We defined “secretory velocity” by using scAPEX-seq counts as a proxy for spliced reads, and Supernatant-seq counts as a proxy for unspliced reads. Secretory velocity trajectories were distinct from those of RNA velocity, showing movement toward committed effector and basally-active states, and away from exhausted and memory-like states (**Fig. 5H**). Increasing secretory commitment may thus represent a complementary axis of differentiation that is relevant for engineering CAR T cells towards durable effector phenotypes.

### CTSW improves long-term persistence and function of CAR T cells

Memory-like CAR T cells are associated with durable clinical responses and long-term persistence^47^. We therefore focused on identifying CCI-induced programs that promote memory-like states or sustained effector function. In short-term coculture experiments, scAPEX-seq revealed robust upregulation of NT5E (also known as CD73) in memory CAR T cells (**Fig. 4I**). Notably, scAPEX-seq showed that NT5E upregulation was not limited to transcript abundance, but was accompanied by a pronounced increase in ER localization **(Fig. 4I)**. Given that NT5E catalyzes the production of immunosuppressive adenosine, and that other adenosine-related genes such as SLC29A1 were also upregulated, we hypothesized that NT5E induction following CAR activation contributes to progressive CAR T cell dysfunction. To test this, we first used Dynamo’s learned analytical vector field function to perform *in silico* knockout of NT5E. This predicted enhanced differentiation towards effector function and away from exhaustion, supporting the possibility that NT5E restrains CAR T cell function (**Fig. S6A**).

Experimentally, we confirmed that NT5E protein levels were increased in CC compared to NCC CAR T cells (**Fig. S7A**). Untransduced CD8 T cells also showed less NT5E induction under identical CC conditions (**Figs. S7A-B**). We then generated NT5E knockout CAR T cells using CRISPR (**Fig. S7C**). In a repeat stimulation killing assay, NT5E-KO CAR T cells showed improved killing over time compared to wild-type controls (**Figs. 6A-B**). The exhaustion markers PD1, TIM3, and LAG3 were decreased in these cells (**Fig. S7D**). These findings were recapitulated using a lower-affinity anti-HER2 CAR^48^, suggesting that NT5E deletion can be beneficial under conditions of weak or chronic antigen engagement (**Figs. 6B, S7E**), consistent with recent findings by Klysz et al^49^.

**Figure 6.**
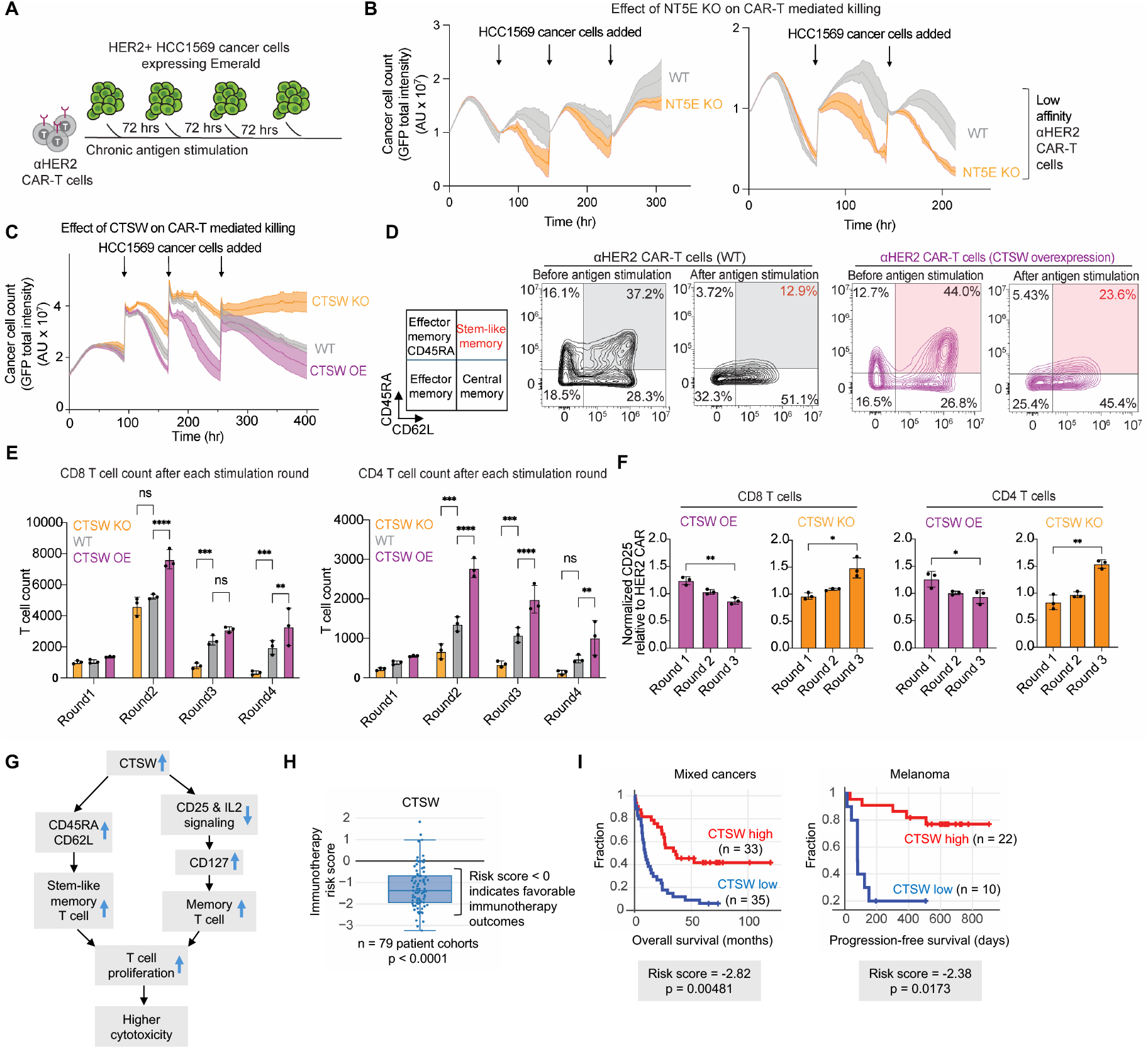
CTSW improves the long-term persistence of CAR T cells. (**A**) Schematic of CAR T cell persistent stimulation assay. Anti-HER2 CAR T cells were co-cultured with HER2^+^ breast cancer cells (HCC1569) expressing Emerald. Every three days, CAR T cells were transferred onto fresh cancer cells. Cancer cell fluorescence intensity is continuously quantified by IncuCyte using expressed Emerald marker. (**B**) Results of persistent antigen stimulation assay in (A) using CD8^+^ anti-HER2 CAR T cells with or without NT5E knocked out (KO). Ratio of CAR T cells to tumor cells at the start of each stimulation cycle was 1:10. Either a high affinity^33^ or low-affinity^48^ anti-HER2 CAR was used. Plots show mean ± SD of three technical replicates. (**C**) Persistent antigen stimulation assay using CD3^+^ anti-HER2 CAR T cells (high affinity) with CTSW knocked out (KO) or overexpressed (OE). Ratio of CAR T cells to tumor cells at the start of stimulation was 1:10. Plots show mean ± SD of three technical replicates. N = 3 donors, with representative data from one donor shown. (**D**) CTSW overexpression increases the fraction of CAR T cells in a stem-like memory state after antigen stimulation, as indicated by CD62L and CD45RA staining. Proportion of CD62L^+^ CD45RA^+^ CAR T cells indicated in top right quadrant. (**E**) CTSW overexpression promotes sustained proliferation of both CD8^+^ (left) and CD4^+^ (right) CAR T cells. Flow cytometry quantification of CAR T cell numbers after each round of cancer cell stimulation. ns = not significant, *p < 0.05, **p < 0.01, ***p < 0.001, ****p < 0.0001 (two-way ANOVA). (**F**) CTSW modulates CD25 expression in CAR T cells. Flow cytometry analysis of surface CD25 expression on CD8^+^ (left) and CD4^+^ (right) anti-HER2 CAR T cells following repeated cancer cell stimulation. *p < 0.05, **p < 0.01 (two-tailed Student’s t test). (**G**) Proposed model for how CTSW promotes CAR T cell cytotoxicity. (**H**) CTSW expression is correlated with improved immunotherapy outcomes. Analysis of bulk transcriptomic datasets using the Cancer Immunology Data Engine (CIDE)^53^ indicates that across 79 patient cohorts with documented responses to immune checkpoint blockade, CTSW expression is inversely correlated with immunotherapy risk score. (**I**) CTSW expression correlates with improved survival in multiple human cancers. Kaplan–Meier survival analyses of patients with non-small cell lung cancers^54^ (left) or melanoma^55^ (right).

While NT5E represents a CCI-induced negative regulator, we sought to identify genes associated with sustained CAR T cell function. From long-term CC scAPEX-seq data, we focused on the late effector CAR T cell cluster that we hypothesized represents a persistent, functional population. DEG analysis highlighted several candidate genes (*IFI6, APOD*, and *CTSW*) that met two criteria: (1) they are upregulated in the late effector cluster relative to early (2 hr) major effector CAR T cells, and (2) they are downregulated or absent in terminally-exhausted CAR T cell populations (**Fig. 5E**). We individually overexpressed each candidate gene in anti-HER2 CD8^+^ CAR T cells to test their functional impact.

Among the candidates, CTSW (Cathepsin W) emerged as the most promising enhancer of CAR T cell persistence. Dynamo analysis predicted that CTSW overexpression would divert CAR T cells away from exhaustion and toward the late effector state (**Fig. S6B**). Consistent with this prediction, CTSW overexpression (OE) offered the most pronounced improvement in long-term tumor cell killing in repeated stimulation assays (**Fig. S8A**). While effects were minimal during the first round of stimulation, CTSW’s benefits increased with subsequent rounds. The same beneficial phenotype of CTSW-OE was observed in a related experiment with human CD3^+^ T cells containing both CD4^+^ and CD8^+^ subsets (**Fig. 6C**). Conversely, CTSW knockout (KO) resulted in failure to control tumors (**Figs. 6C, S8B-D**).

CTSW is a member of the papain-like family of cysteine proteases and is primarily expressed in cytotoxic effector cells such as CD8^+^ T cells and natural killer (NK) cells^50^. A 21-amino acid N-terminal signal peptide directs CTSW to the ER membrane. Although CTSW substrates are incompletely characterized, prior work has shown that CTSW can cleave CD25 (IL2R alpha) on peripheral regulatory T cells^51^. Because reduced CD25 expression is associated with long-lived (high CD62L), stem-like memory states^52^, we hypothesized that this may be one mechanism by which CTSW enhances CAR T cell persistence.

We found that CTSW-overexpressing CAR T cells exhibited an increased fraction of CD62L^+^CD45RA^+^ cells both before and after tumor stimulation, indicative of a stem-like memory phenotype (**Fig. 6D**). Since durable proliferation is a hallmark of stem-like T cells, we next quantified T cell expansion after each round of antigen stimulation. While there were no significant differences in T-cell numbers during the initial round (**Fig. 6E**), CTSW-OE CAR T cells expanded significantly more than wild-type controls from the second round onward, whereas CTSW-KO cells showed impaired proliferation. This proliferative advantage was observed in both CD4^+^ and CD8^+^ subsets, suggesting a broad role for CTSW in sustaining long-term T cell fitness.

In parallel, we found that when compared to unmodified CAR T cells, CTSW OE led to a progressive reduction in CD25 expression across repeated stimulation cycles, whereas CTSW KO produced the opposite effect in both CD4^+^ and CD8^+^ CAR T cells (**Figs. 6F, S8E**). Together, these data support a model in which CTSW promotes durable CAR T cell function by maintaining a CD62L^+^CD45RA^+^ stem-like state and attenuating CD25 expression, thereby enhancing self-renewal and proliferation (**Fig. 6G**).

Finally, to assess the clinical relevance of CTSW expression, we analyzed patient outcomes using the Cancer Immunology Data Engine (CIDE)^53^, which integrates 90 omics datasets spanning 8,575 tumor profiles with documented immunotherapy outcomes. CTSW displayed a significant negative immunotherapy risk score, indicating that higher CTSW levels predict greater therapeutic benefit of immune checkpoint blockade (**Fig. 6H**). In both melanoma and non-small cell lung cancer^54,55^, elevated CTSW expression in the tumor microenvironment (likely in CD8^+^ T cell and NK cell populations) is correlated with improved overall patient survival following immune checkpoint blockade (**Fig. 6I**), consistent with our finding that CTSW overexpression enhances cytotoxic function and persistence of CAR T cells. Taken together, these findings identify CTSW as a T cell–intrinsic regulator of persistence and effector function that could be leveraged to enhance the efficacy of adoptive T cell therapies in solid tumors. Given its expression in NK cells, CTSW may also have broader implications, beyond T cells, for cytotoxic lymphocyte-based immunotherapies.

## Discussion

Proximity labeling has been widely applied to map subcellular proteomes and interactomes by mass spectrometry^56^, but its application to cellular RNAs has been much less extensive, in part because of low RNA capture efficiency^10^. Here, replacement of BP with the more cell permeable PA probe and protocol improvements to increase RNA recovery yielded APEX-seq2, a next-generation bulk spatial transcriptome mapping method with nanometer spatial resolution and 57-100% transcript coverage across multiple compartments, using as little as 4 μg of input RNA.

APEX-seq2 paved the way for the development of single cell APEX-seq. By using streptavidin beads to fractionate APEX-labeled, single cell-barcoded transcriptome libraries prior to amplification, we obtained unbiased subcellular transcriptome profiles from thousands of single cells simultaneously. Single cell technologies are most valuable for the analysis of heterogeneous samples in which a diversity of cell types interact to produce complex behaviors. To gain insight into such cell-cell interactions, we focused on transcripts localized to the ER membrane that encode cell surface and secreted proteins (e.g., ligands and receptors) that mediate the bulk of intercellular communication.

We found that scAPEX-seq resolves distinct cell states and captures coculture-induced changes that are missed by conventional scRNA-seq. One reason is that while single-cell technologies generally suffer from low sensitivity and drop out of genes, scAPEX-seq captures and accurately quantifies transcripts most relevant to the biology of interest. Furthermore, many studies have shown that RNA abundance is poorly correlated with total protein expression, and especially with cell surface protein copy number. In situations where CCIs induce changes in the cell surface/extracellular proteome through mRNA relocalization or changes in RNA accessibility rather than total abundance, scAPEX-seq may be able to capture these changes while whole transcriptome sequencing cannot. Indeed, we observed dozens of immune-relevant transcripts that show altered labeling at the ER membrane induced by coculture in our macrophage-cancer coculture dataset.

Because APEX2 labeling is like a contour map that enriches ER-proximal transcripts most strongly but still produces detectable labeling hundreds of nanometers away, transcripts from other compartments are not completely absent in the scAPEX-seq fraction, just de-enriched. This likely contributes to the number of features and UMIs recovered in scAPEX-seq samples. Although the broader labeling radius is often viewed as a limitation, here it provides useful cellular context for analysis. In addition, we retain unlabeled transcripts for Supernatant-seq (an approximation of whole-transcriptome seq), as a simple byproduct of the scAPEX-seq workflow. These two views of the same cells can enable new methods of multimodal analysis that may further extend the utility of scAPEX-seq.

scAPEX-seq can be compared to other modern technologies for spatial transcriptome analysis. Imaging-based methods such as MERFISH^7^, seqFISH^8^, and STARmap^9^ provide valuable tissue context, but for the analysis of subcellular RNA populations such as the secretome, scAPEX-seq is likely to offer higher transcript coverage, spatial specificity, cell throughput, and ease of use. Slide/array-based RNA capture methods, such as Slide-seq^57^, Slide-tags^58^, Stereo-seq^59^, Seq-scope^60^, and Open-ST^61^ provide higher transcript coverage (UMIs per cell) than imaging methods, and newer techniques achieve single cell or even subcellular resolution by reducing cross-cell capture. However, when compared to scAPEX-seq, these methods are generally limited by RNA capture efficiency, cost, complicated experimental protocols and analysis, and loss of spatial resolution due to RNA diffusion before capture. Other technologies, including CITE-seq^62^ and SEC-seq (using hydrogel nanovials)^63^, can similarly be used for CCI inference through profiling cell surface and secreted proteins with DNA-conjugated antibodies – and will likely be complementary to scAPEX-seq.

An important consideration when using spatial transcriptomic methods is that while spatial context is certainly advantageous in CCI prediction, spatial proximity of two cells does not equate to interaction, and inference of CCIs through other means is still required. Analytical methods like CellPhoneDB^64^, CellChat^65^, and NicheNet^66^ map co-expression patterns of known ligand-receptor pairs to identify cell clusters that are likely to be in communication; these have shown a strong ability to uncover biologically important CCIs. We also used such methods of analysis for scAPEX-seq data, and found that more interacting clusters and unique ligand-receptor pairs could be identified, due to the greater sensitivity and dynamic range of scAPEX-seq for quantifying CCI-relevant transcripts.

We applied scAPEX-seq to study the interactions of cancer cells with both macrophages and CAR T cells. For CAR T cells, scAPEX-seq uncovered previously unresolved subclusters with distinct secretory transcriptomes and cell state trajectories as inferred by Dynamo^46^. Among early coculture induced programs, we identified NT5E (also known as CD73), a key mediator of immunosuppressive adenosine within the tumor microenvironment^37^. Genetic ablation of NT5E reduced exhaustion markers, enhancing T cell function and cytotoxicity. Notably, scAPEX-seq revealed early engagement of adenosine-mediated immunoregulatory pathways in memory CAR T cells within just two hours of tumor contact, suggesting that inhibitory and modulatory programs are rapidly initiated during CAR T cell engagement. These early transcriptional and localization-dependent signals may represent actionable targets to modulate CAR T cell fate before exhaustion becomes established.

We also studied CTSW, a gene that was upregulated in a scAPEX-seq-specific cluster of long-lived CAR T cells with reduced exhaustion markers and improved expression of effector molecules. This cell population was not visible using the whole transcriptome (Supernatant-seq). Recently^67^, CTSW was independently suggested as an important immunomodulatory molecule in CAR T cells, but through multi-omics approaches and with a specific focus on cysteine proteases. Our work demonstrates, using repeat antigen exposure assays, that CTSW expression contributes to long-term cancer cell killing, is associated with sustained CAR T persistence, and supports a stem-like phenotype. These findings highlight how subcellular transcriptomic profiling can reveal regulators of immune cell fate that remain hidden in RNA abundance-only datasets.

Beyond immuno-oncology, cell-cell interactions are critical to a vast array of other processes that underlie the biology of multicellular organisms, including tissue development, aging, pathogen infection, and cognition. scAPEX-seq addresses a fundamental limitation of conventional scRNA-seq in CCI inference, which treats total mRNA abundance as a proxy for protein output and interaction potential. By selectively enriching ER-localized secretomes, scAPEX-seq refines interaction predictions and reveals functionally important cell states and regulatory pathways that are difficult to resolve at the level of total transcript abundance alone. While this study focused on the ER membrane due to its relevance to CCIs, scAPEX-seq should be easily extensible to other important subcellular regions, such as the mitochondria, stress granules, and synapses, enabling systematic investigation of how RNA localization and dynamics shape complex cell and tissue behaviors.

## Acknowledgements

A.Y.T. is grateful to the National Science Foundation (MFB Grant 2330686), the CIRM Center for Neuropsychiatric Stem Cell Proteomics, and the Chan Zuckerberg Biohub – San Francisco for funding. R.A.H.-L. holds a Career Award at the Scientific Interface from Burroughs Welcome Fund. A.Y.T. and R.A.H.-L. are both Biohub, San Francisco investigators and R.A.H.-L. is a Stanford member researcher of the Parker Institute for Cancer Immunotherapy. A.G.X. is supported by an NSF Graduate Research Fellowship and NIH T32GM136568. Xiaojie Qiu is grateful for the NIH Director’s New Innovator award (1DP2HG014282-01) and NIH’s pathway to independence award (5R00HG012887-03). Nianping Liu is supported by NHGRI MorPhiC Initiative (U01 HG013176). We thank Shizhong Dai and Joleen Cheah for help with stable cell line generation. We thank Howard Chang, Julia Belk, Sifei Yin, Nicholas Kalogriopoulos and Matthew Ravalin for helpful discussions, advice, and analyses.

## Author contributions

A.Y.T. conceived the project. B.C., A.G.X., R.A.H.-L. and A.Y.T. designed the research. B.C. and A.G.X. developed bulk APEX-seq2, scAPEX-seq, and performed scRNA-seq on macrophage–tumor cell cocultures. Q.X., B.C. and R.A.H.-L. designed tumor-CAR T cell coculture experiments, which were performed by B.C. and Q.X. B.C. and A.G.X. prepared sequencing libraries. A.G.X. analyzed all sequencing data with assistance from N.P.L. on tumor-CAR T cell coculture data. N.P.L., X.J.Q. and A.G.X. used Dynamo to analyze cell state transitions. Q.X. and B.C. performed validation experiments on NT5E and CTSW. Q.X. performed repeat antigen stimulation assays and flow cytometry to validate NT5E/CTSW results. A.Y.T. and R.A.H.-L. provided supervision. A.Y.T., B.C., A.G.X., and R.A.H.-L. wrote the paper with input from all authors.

## Declaration of interests

The authors have no competing interests to declare.

## Notes

### Competing Interest Statement

The authors have declared no competing interest.

