## Supplementary information for "Subcellular transcriptome sequencing with single cell APEX-seq identifies regulators of cell-cell interactions"

for

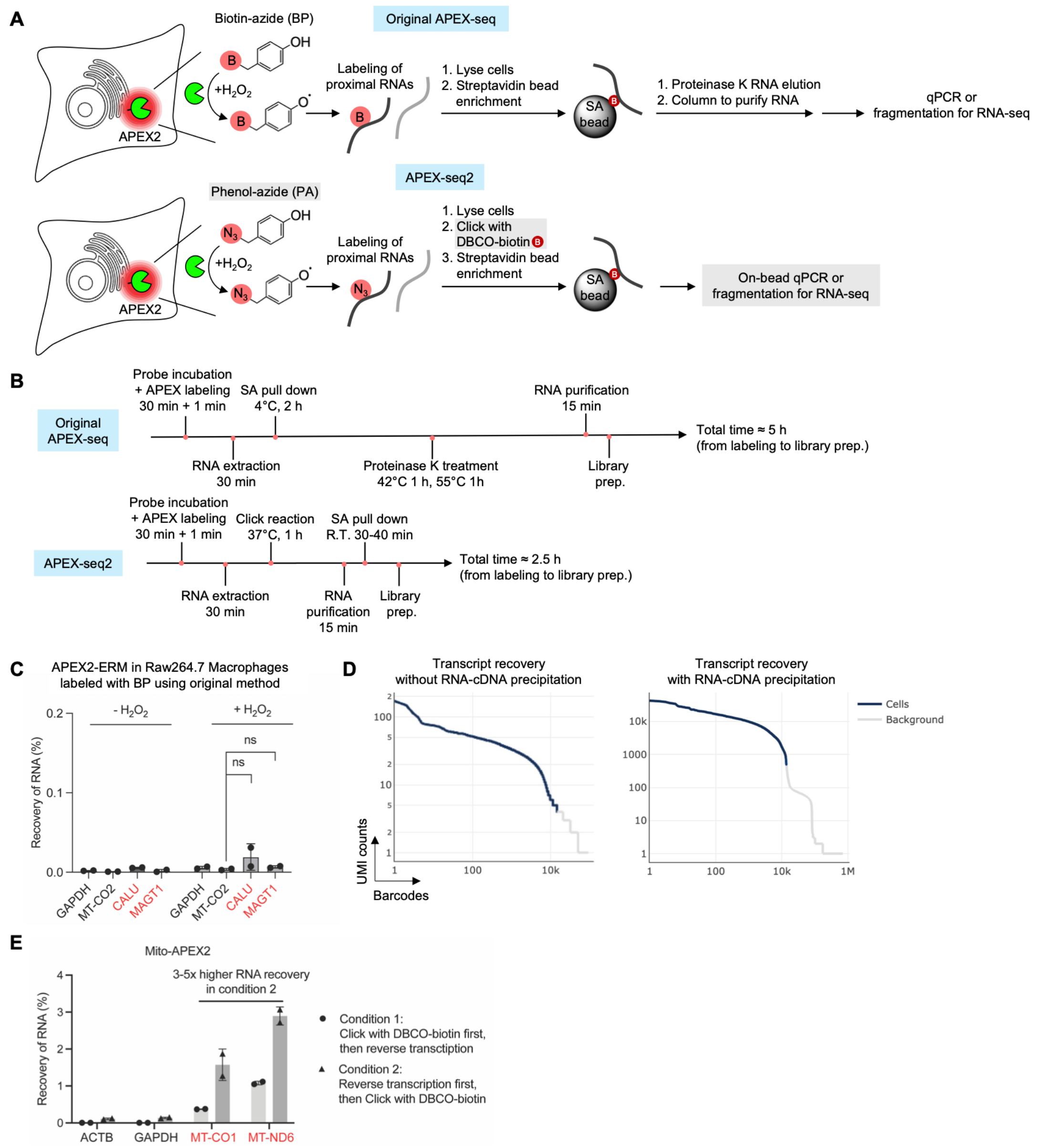


**Figure S1. Optimization of APEX-seq2.** Related to Figure 1. (**A**) Schematics showing original APEX-seq and new APEX-seq2 reported here. (**B**) Workflow timelines for APEX-seq and APEX-seq2. (**C**) RNA recovery using original APEX-seq and BP in Raw264.7 cells. Cells expressed APEX2-ERM and were labeled with 500 μM BP for 1 minute. On-target secretory transcripts (red) and off-target cytosolic and mitochondrial transcripts (black) were quantified by qRT-PCR before and after streptavidin enrichment, to determine fraction of RNA recovered. n = 1 experiment, 2 replicates per condition. Data are mean ± s.d. P-values were determined using two-tailed Student's t tests. (**D**) RNA-cDNA precipitation increases recovery of transcripts and UMI counts by ~100-fold. Cell barcode x UMI count plot for a scAPEX-seq sample without precipitation of RNA (left) or with GlycoBlue-assisted precipitation of RNA-cDNA (right). Raw264.7 macrophages expressing APEX2-ERM were used. (**E**) RNA recovery using APEX-seq2 when reverse transcription is performed before vs after Click with DBCO-biotin. HEK293T cells expressing APEX2 targeted to the mitochondrial matrix were labeled with PA for 1 minute. On-target mitochondrial (red) and off-target non-mitochondrial (black) transcripts were quantified by qRT-PCR before and after streptavidin enrichment, to determine fraction of RNA recovered. n = 2 experiments, 2 replicates per condition. Data are mean ± s.d.

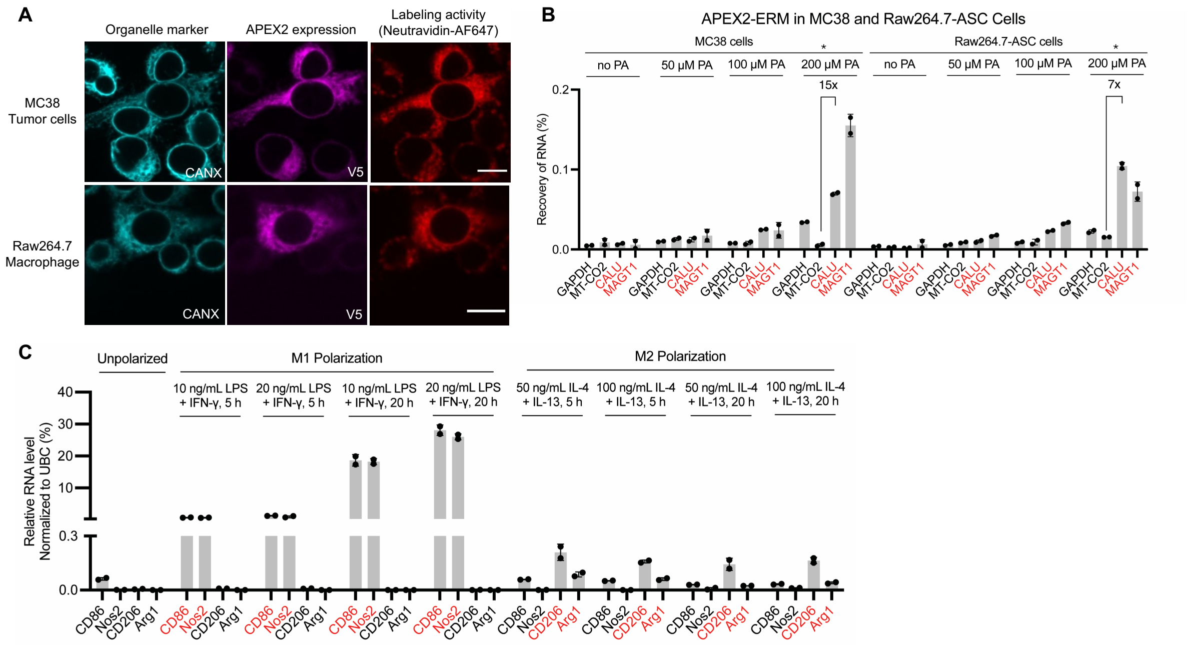


**Figure S2. Characterization of APEX2-ERM-expressing macrophage and MC38 cancer cells.** Related to Figure 2. (**A**) Fluorescence imaging of MC38 tumor cells and Raw264.7 macrophages stably expressing ERM-APEX2. BP labeling was performed live for 1 minute, then cells were fixed and stained with anti-calnexin antibody (CANX, cyan) to visualize the ER, anti-V5 antibody to detect APEX expression (magenta), and neutravidin-AF647 (red) to detect biotinylation. Scale bars, 10 μm. (**B**) Optimization of PA probe concentration for labeling of macrophage-cancer cell cocultures. After 1 minute labeling at the indicated concentrations, on-target ER (red) and off-target (black) transcripts were quantified by qRT-PCR before and after streptavidin enrichment. MT-CO2 is cytochrome c oxidase II and CALU/MAGT1 are ER-associated genes. *indicates best condition, used in experiments in **Fig. 3B**. n = 2 experiments, 2 replicates per condition. Data are mean ± s.d. (**C**) Optimization of M1/M2 polarization conditions for Raw264.7 macrophages cells. qRT-PCR detection of CD86 and Nos2 (M1 markers) and CD206 and Arg1 (M2 markers). Ubiquitin C (UBC) is a housekeeping gene. n = 2 experiments, 2 replicates per condition. Data are mean ± s.d.

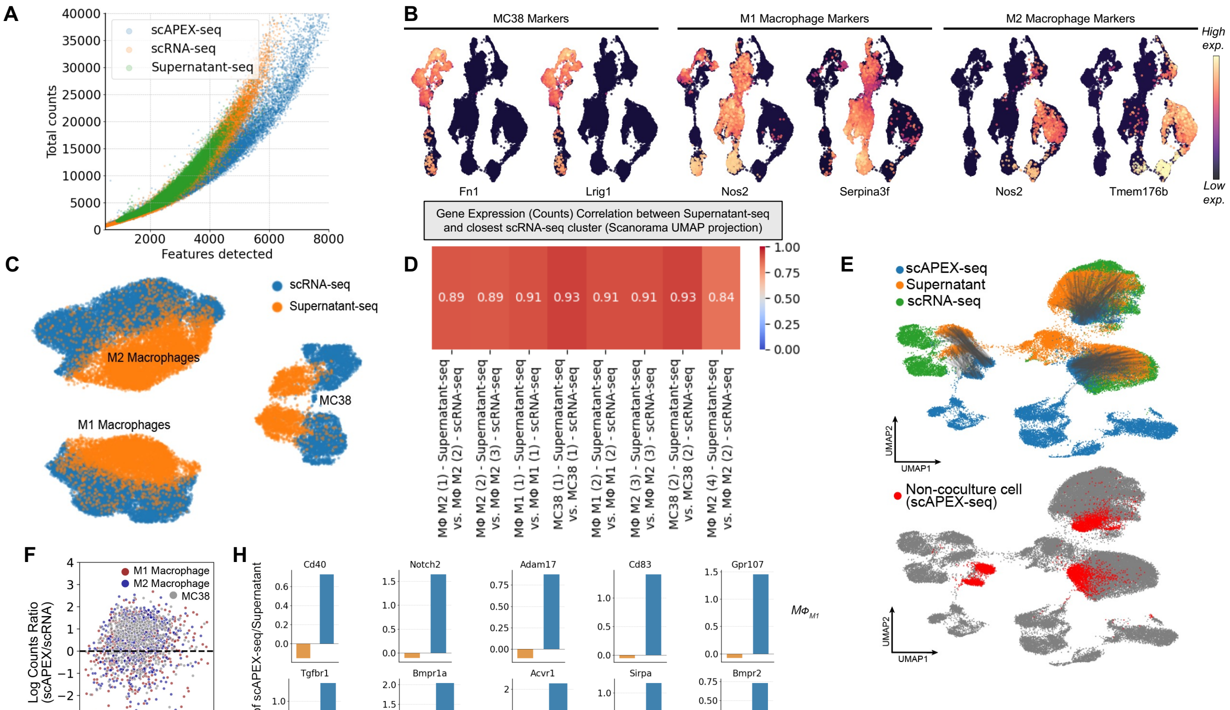


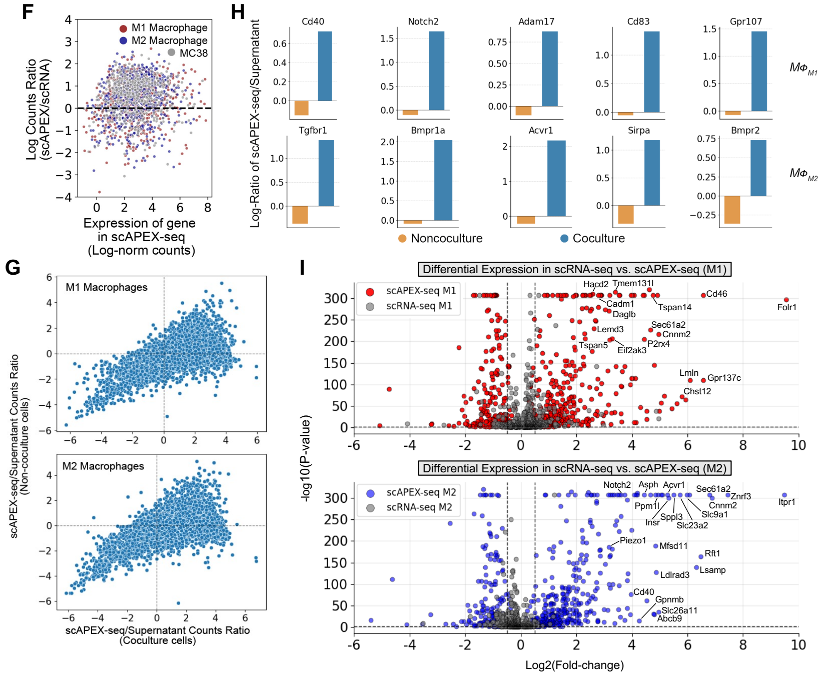


J

| M1 Gene(s) | Relevant Function |
|---|---|
| Cd46 | Complement regulatory protein |
| Daglb | Lipase, regulator of inflammatory lipid signaling |
| Eif2ak3 | ER stress sensor, upstream regulator of macrophage polarization and inflammatory signaling |
| P2rx4 | ATP-gated ion channel, related to inflammasome activation and IL-1β secretion |
| Il6st | Subunit of IL6 receptor complex, regulator of inflammation |
| Lemd3 | Regulator of TGF-β and BMP signaling |
| Lmtk2 | Regulator of NF-kβ activation |

| M2 Gene(s) | Function |
|---|---|
| Itpr1 | Calcium channel, regulator of TNF-α signaling |
| Slc9a1 | Transporter, regulator of pH, associated with increased TAM populations |
| Notch2 | Notch signaling promotes M2-like activation |
| Insr | Insulin receptor, impacts macrophage polarization |
| Acvr1 | BMP receptor, related to SMAD signaling and NF-kβ signaling |
| Ppm1l | Phosphatase, regulator of inflammatory cytokine production |
| Piezo1 | Mechanosensor, stiffness-dependent anti-inflammatory activity |
| Gpnmb | Transmembrane glycoprotein, negative regulator of inflammation in macrophages |
| Cd40 | Costimulatory receptor, inflammation promoting |

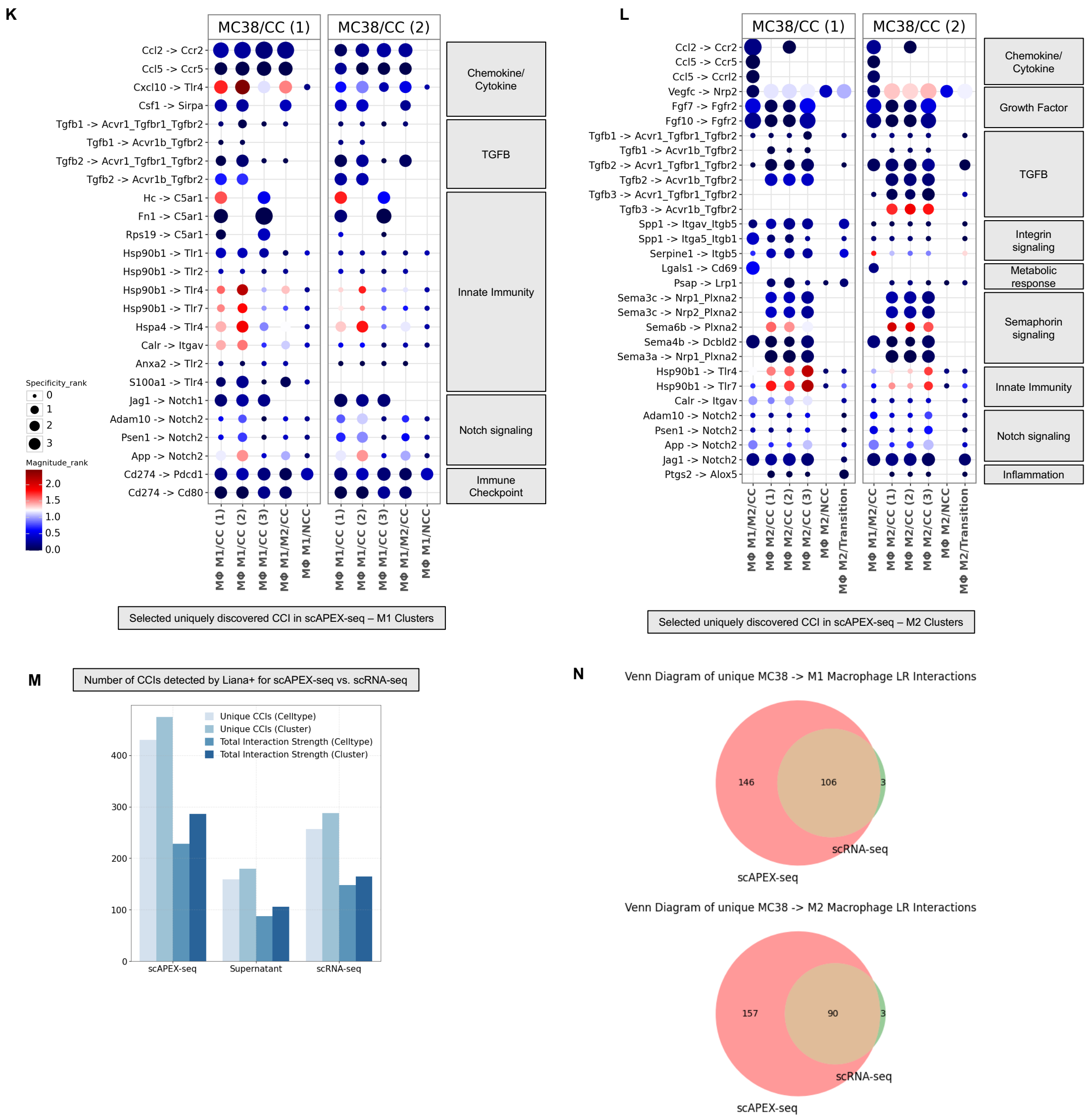


**Figure S3. Additional data on macrophage-cancer cell cocultures.** Related to Figure 3. (**A**) Scatter plot showing total counts and total features per cell, for scRNA-seq, Supernatant-seq, and scAPEX-seq. (**B**) scAPEX-seq UMAP plots highlighting cell type-specific markers for MC38, and M1/M2 macrophages. (**C**) Scanorama integration of Supernatant-seq and scRNA-seq-identified cell clusters, colored by sample of origin. (**D**) Heatmap showing gene expression correlation (spearman) between cell clusters identified in Supernatant-seq and the closest matched cluster in scRNA-seq (as defined by cluster centroid distance on UMAP). (**E**) Integration of scRNA-seq, Supernatant-seq, and scAPEX-seq data by Scanorama, colored by sample of origin (top) or NCC cells (bottom). (**F**) Scatter plot showing scAPEX-seq sensitivity for low-count secretory genes. Log2 fold change of scAPEX-seq counts over scRNA-seq counts for each gene is shown, separated by cell type. Line at Log2FC = 0 represents equal sensitivity. Related to **Fig. 3G,** but x-axis is plotted as expression in scAPEX-seq. (**G**) Analysis of gene re-localization upon coculture using scAPEX-seq and Supernatant data. Scatter plots of scAPEX-seq/Supernatant count ratio for CC vs. NCC samples shown for both M1 and M2 macrophage populations. (**H**) Selected genes from **Fig. S3G** with immunomodulatory effects that show increased labeling by APEX2-ERM upon co-culture. (**I**) Summary of DEGs identified between scAPEX-seq and scRNA-seq. Volcano plots show the relative distribution of DEGs, with top hits annotated for scAPEX-seq in the top right. (**J**) Tables show functional annotation of DEGs uniquely identified by scAPEX-seq as upregulated in CC M1/M2 macrophages. (**K-L**) Subset of unique CCIs identified by LIANA+ for M1 and M2 macrophages in scAPEX-seq. Cytokine/chemokine, ECM, and immunosuppressive pathways are featured. (**M**) Bar chart depicting unique interactions discovered by scAPEX-seq, Supernatant-seq, and scRNA-seq when classified by celltype only or by individual clusters. Total interaction strengths as calculated by LIANA+ are also summarized. (**N**) Venn diagrams showing overlap of LIANA+-identified ligand-receptor (LR) interactions between MC38 and M1/M2 macrophages in scAPEX-seq vs. scRNA-seq.

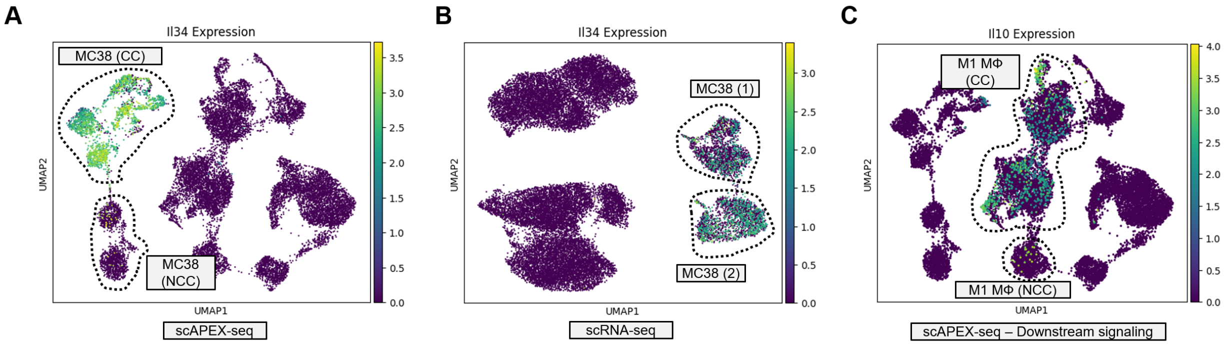


**Figure S4. Analysis of IL34-CSF1R ligand-receptor pair.** (**A-C**) Identification of IL34-CSF1R signaling in scAPEX-seq. Comparison to scRNA-seq is shown in (B), and increased downstream IL10 signaling in M1 macrophages in scAPEX-seq is highlighted in (C).

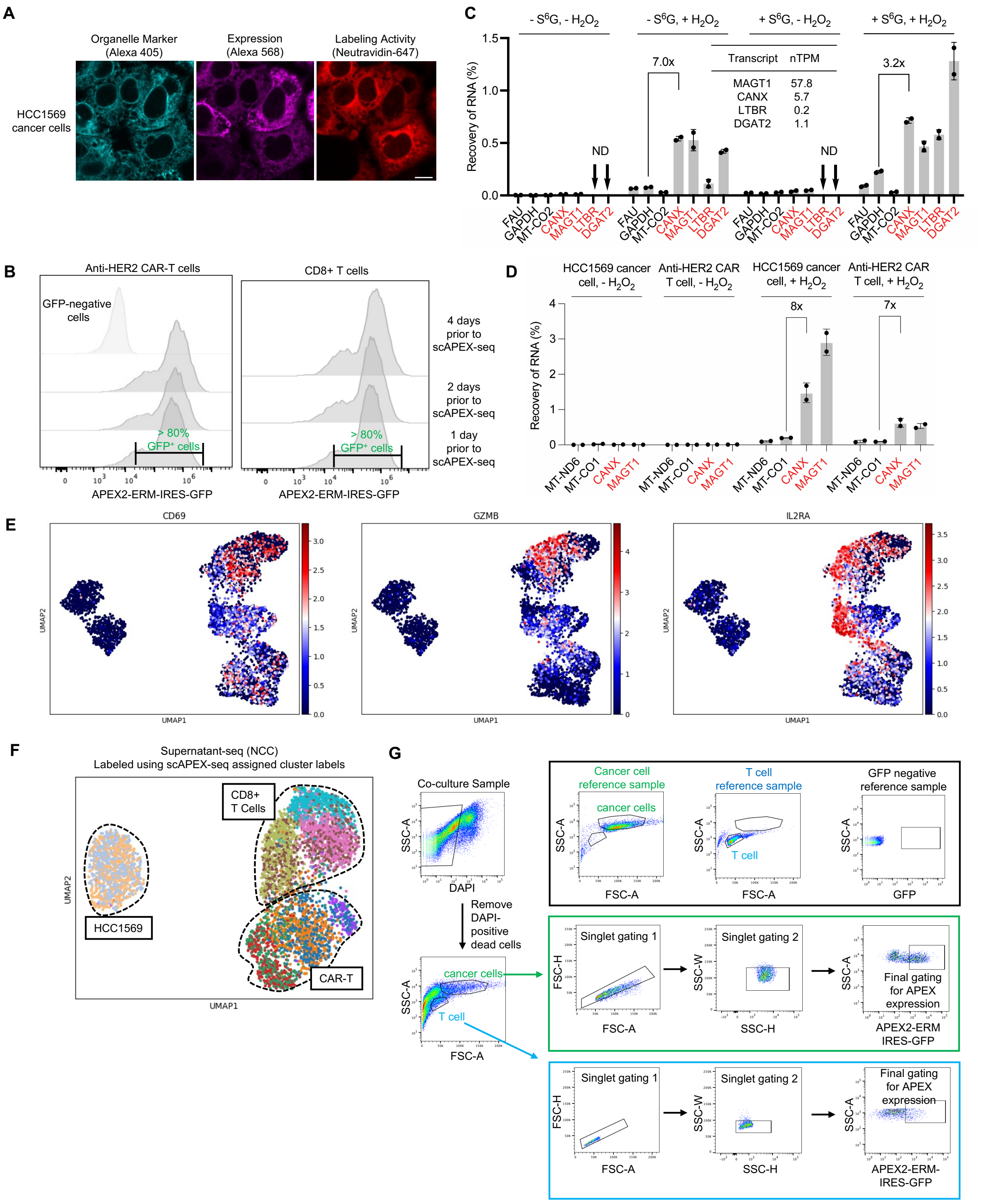


**Figure S5. Additional data on CAR T cell scAPEX-seq.** Related to Figures 4 and 5. (**A**) Fluorescence imaging of APEX2 localization and biotinylation activity at ER membrane (facing cytosol) of HCC1569 cancer cells stably expressing APEX2. Live cell labeling was performed for 1 minute using biotin-phenol, then cells were fixed and stained with anti-Calnexin (cyan) to visualize the ER, anti-V5 to detect APEX expression (magenta), and neutravidin-Alexa Fluor 647 (red) to detect biotinylation. Scale bars, 10 µm. (**B**) Characterization of APEX2 expression in primary anti-HER2 CAR T and CD8$^{+}$ T cells. Transduced cells were sorted, and GFP expression was monitored over 10 days prior to scAPEX-seq. More than 80% of cells remained GFP-positive one day before the experiment. (**C**) $S^6G$ metabolic labeling improves transcript recovery. Bar graph showing the relative recovery of secretory and non-secretory RNAs. HEK293T cells stably expressing ERM-targeted APEX2 were labeled with PA for 1 minute, with or without prior 4-hour treatment with 100 μM 6-thioguanosine ($S^6G$). On-target (red) and off-target (black) transcript levels before and after streptavidin enrichment were quantified by qRT-PCR to determine fraction of RNA recovered. n = 2 experiments, 2 replicates per condition. Data are mean ± s.d. nTPM, normalized transcripts per million. Even low abundance transcripts like DGAT2 are effectively enriched by APEX2. (**D**) $S^6G$ metabolic labeling in CAR T and HCC1569 cancer cells, each stably expressing ERM-targeted APEX2. Bar graph showing the recovery of on-target secretory transcripts (red) and off-target transcripts (black) after 2 hr incubation with $S^6G$ and 1 minute PA labeling. Transcripts were quantified before and after streptavidin enrichment by qRT-PCR to determine fraction of RNA recovered. n = 2 experiments, 2 replicates per condition. Data are mean ± s.d. (**E**) UMAP plots of CAR T NCC scAPEX-seq sample, showing expression of CD69, GZMB, and IL2RA (markers of T-cell activation). (**F**) UMAP plot of CAR T NCC Supernatant-seq sample, showing cell clusters assigned by scAPEX-seq counts for the same cells. Cluster labels are mixed between Supernatant-seq clusters. (**G**) Gating strategy for sorting HCC1569 cancer and anti-HER2 CAR T cells from repeated stimulation assay on Day 27 for scAPEX-seq. DAPI staining was used to remove dead cells.

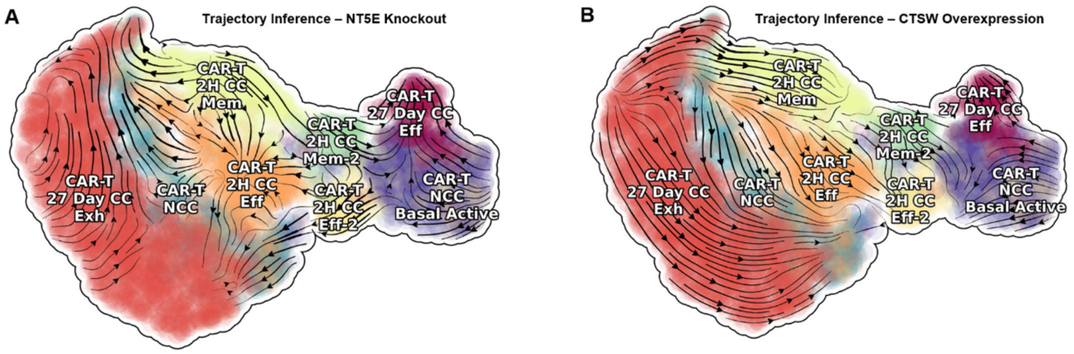


**Figure S6. Dynamo *in silico* perturbation of NT5E and CTSW.** (**A**) UMAP plot of integrated early and late CAR T cells with RNA velocity plotted for NT5E knockout. (**B**) UMAP plot of integrated early and late CAR T cells with RNA velocity plotted for CTSW overexpression.

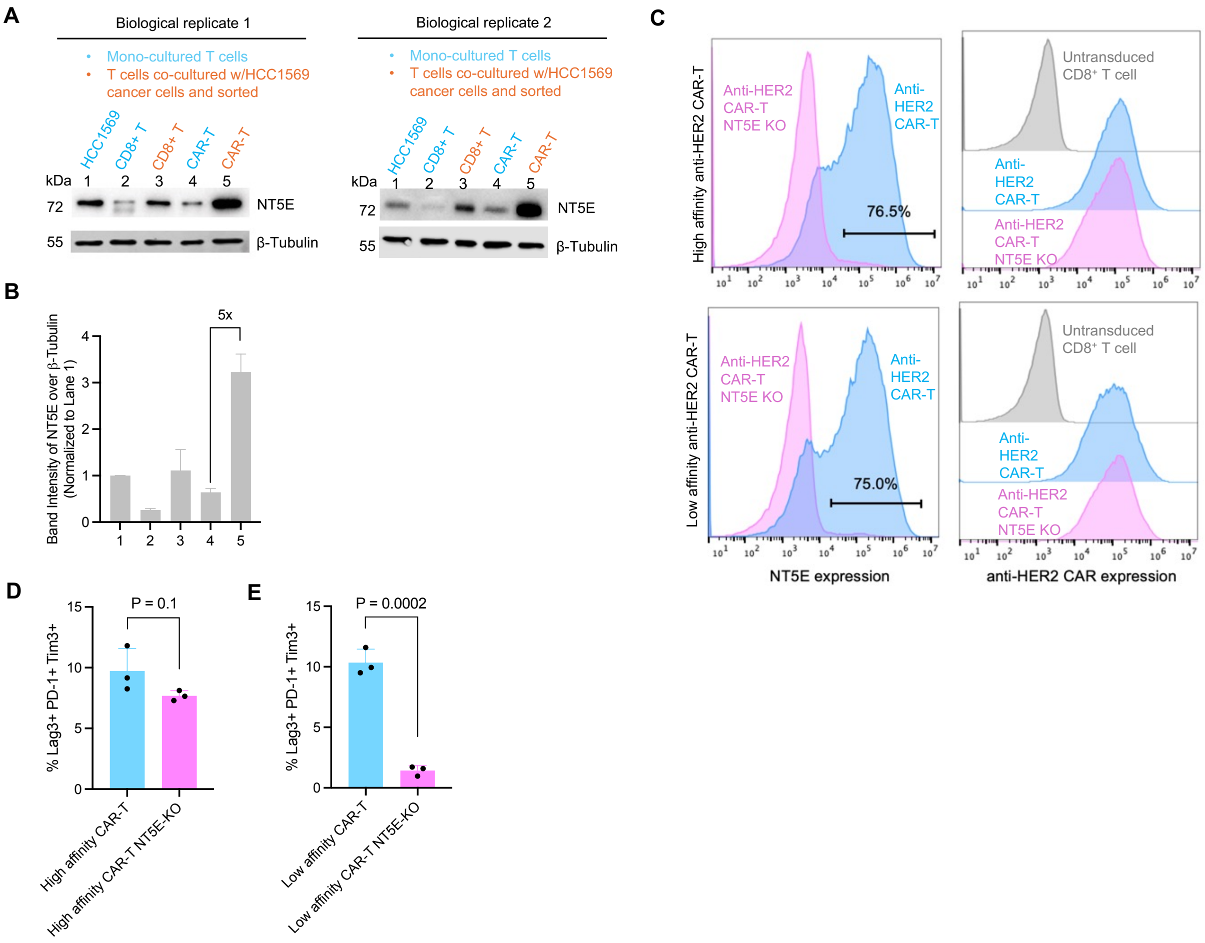


**Figure S7. Additional data on NT5E.** (**A**) Immunoblot analysis of NT5E protein expression levels in HCC1569 cancer cells (lane 1), monocultured CD8+ T cells (lane 2), cocultured $CD8^+$ CAR T cells (lane 3), monocultured high-affinity CAR T cells (lane 4) and cocultured high-affinity CAR T cells (lane 5). For cocultured samples, T cells were separated from cancer cells by sorting prior to lysis and immunoblot analysis. β tubulin was used as a loading control. Two independent biological replicates are shown. (**B**) Quantification of NT5E protein expression from (A). Band intensities were quantified by ImageJ and normalized to corresponding β-tubulin for each lane, and reported relative to HCC1569 cancer cells. Data are mean ± SD from the two replicates. (**C**) Flow cytometry analysis of NT5E expression level in anti-HER2 CAR-$CD8^+$ T cells. Live-cell staining was performed using Anti-NT5E-PE-Cyanine7 to detect NT5E at cell surface. (**D-E**) Proportion of control and NT5E-KO high affinity (**D**) or low affinity (**E**) CAR T cells expressing three exhaustion markers (TIM3, PD1, and LAG3) after continuous rounds of antigen stimulation. Plots show mean ± SD of three technical replicates. N = 3 donors, with representative data from one donor shown.

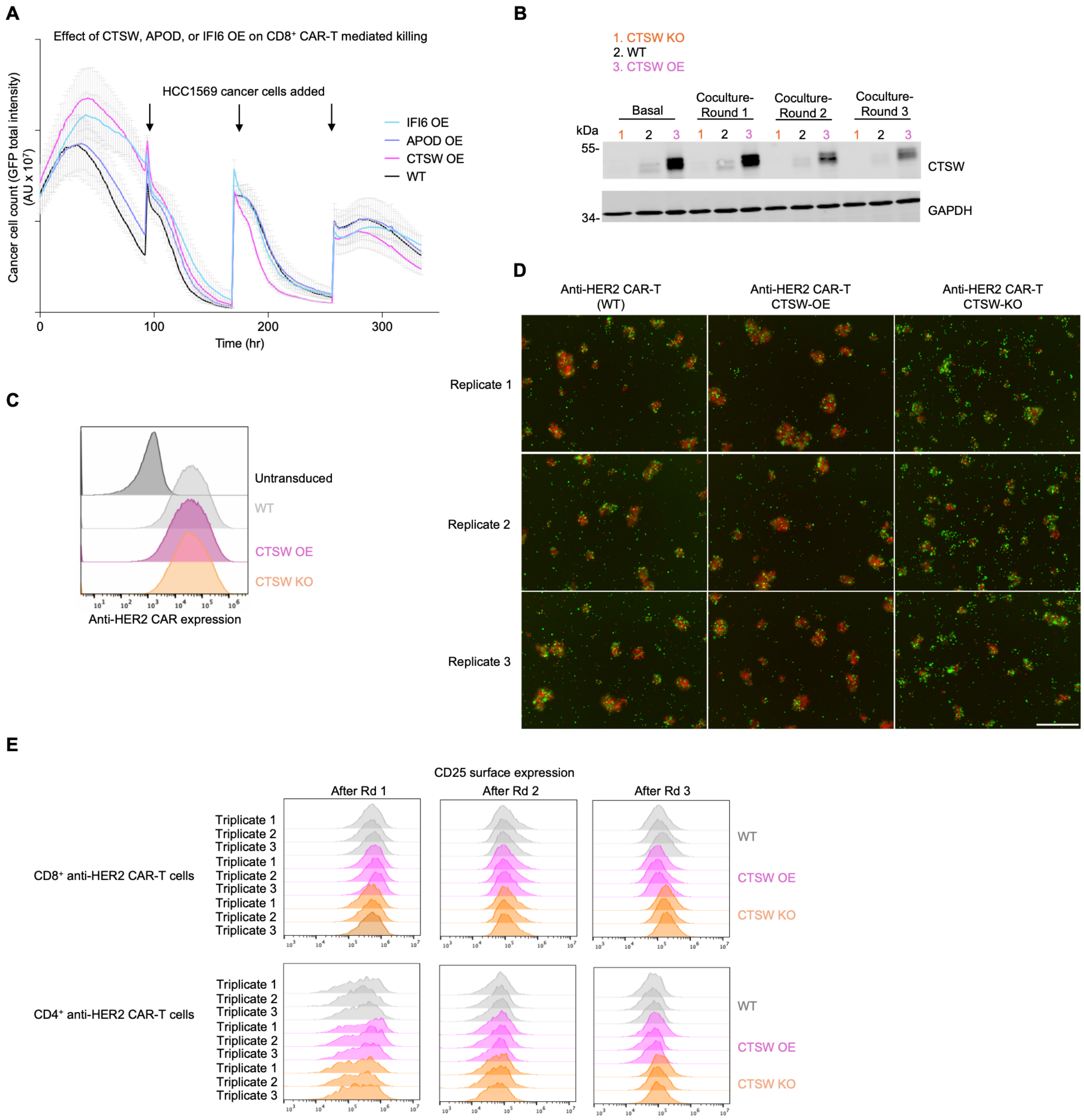


**Figure S8. Additional data on CTSW.** (**A**) Results of persistent antigen stimulation assay in **Fig.6A** using wild-type (WT), CTSW-overexpressing (CTSW-OE), APOD-OE, or IFI6-OE CD8$^{+}$ anti-HER2 CAR T cells. Ratio of CAR T cells to tumor cells at the start of each stimulation cycle was 1:10. Data represent three independent replicates per condition. Error bars represent mean ± SD. (**B**) Immunoblot analysis assessing CTSW expression in wild-type (WT), CTSW-knockout (CTSW-KO), and CTSW-overexpressing (CTSW-OE) CD3$^{+}$ anti-HER2 CAR T cells during persistent antigen stimulation. (**C**) Flow cytometry histograms of anti-HER2 CAR expression level in untransduced control, wild-type, CTSW-KO, and CTSW-OE CD3$^{+}$ CAR T cells. (**D**) Representative fluorescence images from IncuCyte showing coculture of HCC1569 cancer cells (expressing Emerald-H2B (green) in the nucleus) with WT, CTSW-OE, and CTSW-KO CD3$^{+}$ CAR T cells (expressing mCherry marker). Images were acquired at 32 h during the second round of coculture. Scale bars, 400 μm. (**E**) Raw flow cytometry histograms of CD25 surface expression (used for graphs in **Figure 6F**) in CD8$^{+}$ (top row) and CD4$^{+}$ (bottom row) anti-HER2 CAR T cells derived from human primary CD3$^{+}$ T cells.

WT, CTSW-OE and CTSW-KO CAR T cells were analyzed across three sequential rounds (R1–R3) of stimulation with HCC1569 cancer cells.

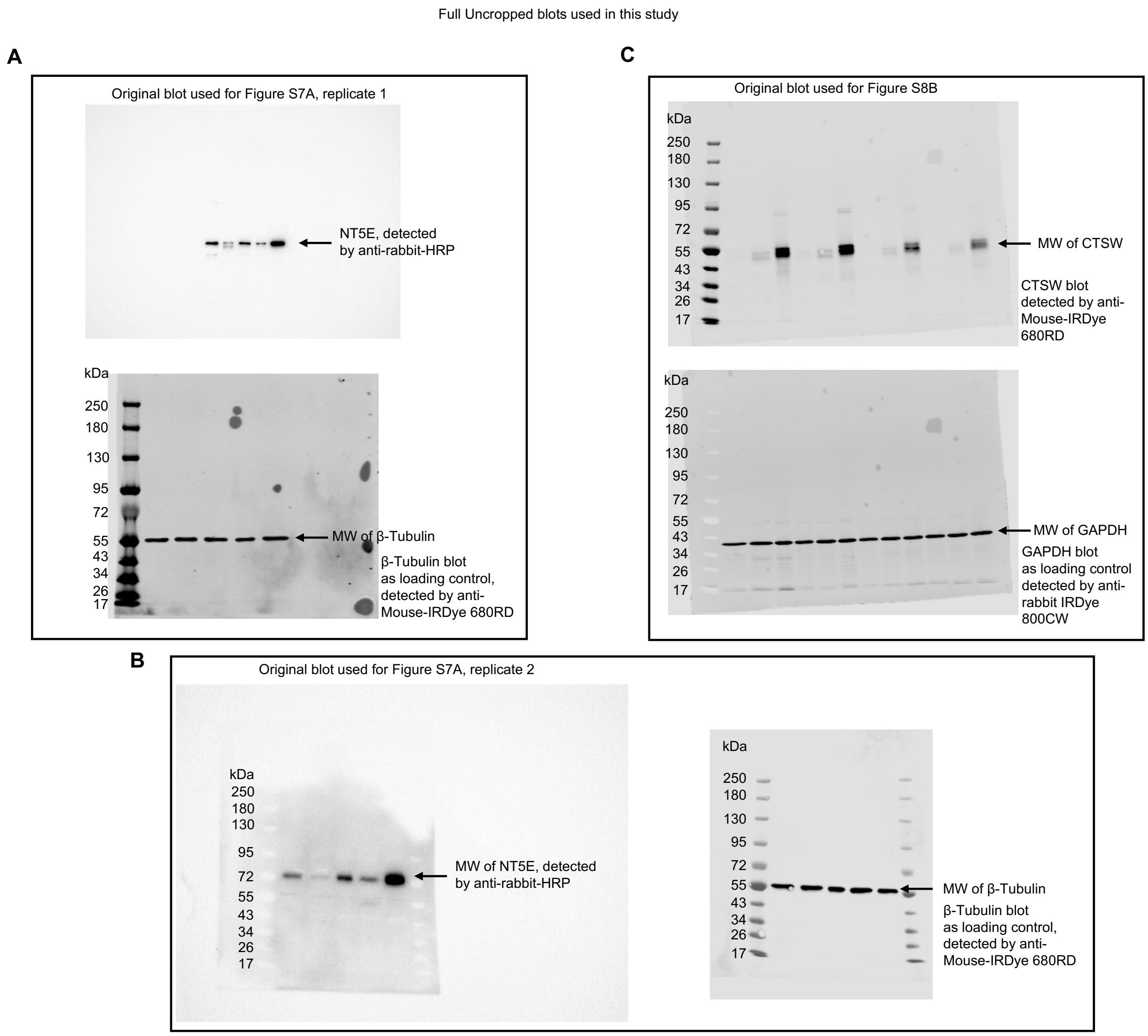


**Figure S9. Full uncropped Western blots used in this study**. (**A-B**) Blot for **Figure S7A.** (**C**) Blot for **Figure S8B.**

### Methods

#### Cloning

Constructs expressing APEX2 with ERM-targeting sequence were generated using standard cloning techniques. PCR fragments were amplified from ERM-APEX2 (Addgene, Cat# 79055) using Q5 polymerase (NEB Cat# M0491S) and cloned into the plx208 lentiviral vector using Gibson assembly. Plasmids were introduced into competent NEB Stable *E. coli* (NEB, Cat # C3040H) via heat shock transformation.

Anti-HER2 CAR constructs were generated by fusing an anti-HER2 single-chain variable fragment (scFv) to the human CD8α hinge domain, followed by the intracellular signaling domains of 4-1BB and CD3ζ. Two variants of anti-HER2 scFv (4D5) were used: a high affinity variant ($K_d$=17.6 nM), addgene, Cat# 164825)[1], and a low affinity variant, 4D5-3 ($K_d$ = 3910 nM)[2]. All CAR constructs included an N-terminal CD8α signaling peptide (MALPVTALLLPLALLLHAARP) to direct membrane localization and a Myc-tag (EQKLISEEDL) for detection of surface CAR expression level.

To generate constructs for lentiviral expression in human primary T cells, the pHR_SFFV lentiviral backbone (Addgene, Cat# 79121) was linearized using BamHI restriction digestion. CAR gene fragments were synthesized by Twist Bioscience and cloned into the digested backbone using In-Fusion cloning (Takara, Cat# 638949). CTSW, APOD and IFI6 overexpression constructs were generated by synthesizing each gene's coding sequence upstream of a T2A-mcherry cassette, then cloned into the linearlized pHR_SFFV lentiviral backbone using the same cloning method mentioned above. Fluorescent reporter constructs encoding H2B-mEmerald (nuclear labeling) and mCherry (cytoplasmitc labeling) were similarly assembled by fusing synthesized fragments into the linearized pHR_SFFV backbone. Plasmids were introduced into Stellar competent *E. coli* (Takara, Cat# 636766 via heat shock transformation.

#### Mammalian cell culture

HEK293T cells (ATCC, Cat# CRL-3216) MC-38 (Kerafast, Cat# ENH204-FP) and Raw264.7-ASC cells (Invivogen, Cat# raw-asc) were cultured in DMEM (Dulbecco's modified Eagle's medium, Gibco Cat# 11965-092) supplemented with 10% (w/v) FBS (fetal bovine serum, VWR Cat# 97068-085) and 1% penicillin-streptomycin (VWR Cat# 16777-164)) at 37 °C under 5% $CO_2$. HCC1569 cancer cells (ATCC, Cat# CRL-2330) were cultured in RPMI 1640 medium (Gibco, Cat# 11875135) supplemented with 10% FBS (Cytiva, Cat# SH30910.03HI) at 37 °C under 5% $CO_2$.

Stable cell lines were generated though lentiviral transduction. To generate lentivirus, HEK293T cells plated at approximately 60% confluency in a 6-well dish were transfected with 2100 ng of the plasmid of interest, 1500 ng of psPAX2, and 660 ng of pMD2G in 500 μL of Opti-MEM (Thermo Fisher Scientific, Cat# 31985062) with 13 μL of TransIT-LT1 (Mirus Bio Cat# MIR 2300). After 30 h, the supernatant containing the lentivirus was then harvested and filtered through a 0.45 μm PES filter (Thermo Fisher Scientific Cat# 725-2545). 2 mL of the fresh supernatant containing 10 μg/mL polybrene (MilliporeSigma Cat# TR1003G) was then added to HEK293T, MC38 or Raw264.7 cells cultured in each well of a 6-well plate with approximately 50% confluency. After 30 h, cells were lifted and replated on a T25 flask and selected with 1 μg/mL of puromycin for 1 week. Cells were split and expanded when they reached 80% confluency. Cells were maintained under this puromycin selection until the time of experiments. Following transduction, cells were sorted using a BD FACSAria II cell sorter (BD Biosciences) at Stanford Shared FACS Facility (RRID: SCR_017788) to isolate populations with stable and uniform genomic integration of either ERM-APEX2-IRES-GFP or H2B-mEmerald.

**Primary human T cell isolation and culture**

Primary human CD8+ T cells were isolated from leukopaks obtained from anonymous healthy donors via negative selection (STEMCELL Technologies, Cat#15063). Isolated T cells were cryopreserved in CELLBANKER 1 (Zenoaq, Cat# 11910). For all experiments, T cells were cultured in human T cell medium (HTCM) composed of X-VIVO 15 (Lonza, Cat# 04-418Q) supplemented with 5% human serum (Bio IVT, HP1022HI), 10 mM neutralized N-acetyl L-cysteine (Sigma-Aldrich, Cat# A9165), and 30 U/mL recombinant human IL-2 (NCI BRB Preclinical Repository).

**Lentiviral transduction of human T cells**

Lentivirus was generated by co-transfecting Lenti-X 293T cells (Takara, Cat# 632180) with three plasmids: the packaging plasmid pPCVd8.91, the envelope plasmid pMD2.G and transgene plasmid. Plasmids were mixed and transfected using PEI-MAX (Polysciences, Cat# 24765). Primary human CD8+ T cells were thawed and rested for 24 hours before stimulation with human anti-CD3/CD28 Dynabeads (Life Technologies, Cat# 11131D) at a 1:1 bead-to-cell ratio. After 48 hours of activation, viral supernatant was harvested, filtered through 0.45 μm filters, and added directly to the T cell cultures for 24 hours. On day 5 post-stimulation, Dynabeads were removed.

**Human T cell sorting and expansion**

T cells were stained using an anti-Myc antibody (Cell Signaling Technology, Cat# 2233S) and sorted using a BD FACSAria II cell sorter (BD Biosciences) at Stanford Shared FACS Facility (RRID: SCR_017788) to enrich for uniform transgene expression. Untransduced cells were used as a negative control. Sorted T cells were expanded for at least 9 days and maintained at $0.5 \times 10^6$ cells/mL during resting. Cells were subsequently used for co-culture assays for scAPEX-seq and cytotoxicity assays for validation studies.

**APEX labeling in living cells**

Prior to all APEX labeling experiments, cells were stably transduced to express APEX-IRES-GFP, with GFP fluorescence serving as a proxy for APEX2 expression levels. As low APEX expression resulted in low RNA recovery, cells with high GFP expression were obtained by selection with puromycin (HEK293T cells) or fluorescence-activated cell sorting (primary T cells, MC38, and Raw 264.7 cells). Correct subcellular localization and APEX2 activity were confirmed by immunofluorescence. Labeling conditions were optimized by titrating the phenol-azide probe to maximize signal-to-noise (recovery of on-target transcripts over off-target transcripts), with final concentrations used between 50-200 μM as indicated.

For labeling experiments with HEK293T cells, cell culture plates were pre-treated with 50 μg/mL HFN (human fibronectin, Millipore Cat# FC010) in DPBS (Gibco Cat# 14190-144) for 30 minutes at 37°C before cell plating. This improves the adherence of HEK293T cells to the plates. 18 h after plating HEK293T cells stably expressing the corresponding APEX2 fusion construct, culture medium was aspirated and replaced with fresh medium containing 500 μM biotin-phenol (APEXbio, Cat# A8011) or 50 μM phenol azide (custom small molecule synthesis by Sai Life Sciences, stored at 500 mM stock concentration in DMSO in -80°C). After 30-minute incubation at 37 °C under 5% CO2, 10x $H_2O_2$ (10 mM, made fresh from stock solution stored at 4°C, Sigma Aldrich, Cat# H1009) was then added to cells to achieve a final concentration of 1 mM, with gentle agitation for exactly 1 min. The labeling reaction was immediately quenched by adding an equal volume of freshly made 2x quencher solution containing 10 mM Trolox (Sigma Aldrich, Cat# 238813-1G), 20 mM sodium ascorbate (Sigma Aldrich Cat# A7631-25G) and 20 mM sodium azide (Sigma Aldrich, Cat# A7631-25G) in DPBS (to prepare the quenching reagents, 1 M sodium ascorbate was made by dissolving 198 mg in 1 mL of water, 1 M sodium azide was made by dissolving 65 mg in 1 mL of water, and 250 mM Trolox was prepared by dissolving 125 mg in 2 mL of DMSO with brief sonication to fully dissolve the powder. The final 2× quencher solution was prepared by combining the appropriate volumes of each stock and bringing the volume to 50 mL with DPBS). Cells were then washed twice with DPBS containing 5 mM Trolox, 10 mM sodium ascorbate and 10 mM sodium azide, and once with DPBS. The cells were then prepared for imaging, RT-qPCR, or bulk RNA-seq experiments as described below. For unlabeled control samples, the protocol was identical, except that the $H_2O_2$ addition step was omitted.

For experiments involve labeling with MC38, RAW264.7, HCC1569, or anti-HER2 CAR-T cells followed by enrichment of labeled RNAs, the cell culture plates were not pre-coated with human fibronectin (HFN), and the probe (phenol azide) concentration was optimized as specified. The 30 min-pre-incubation with probe, 1 min labeling with 1 mM $H_2O_2$ and quenching steps remained the same. For imaging experiments with MC38, RAW264.7 and HCC1569 cells, the labeling protocol utilizing BP remained the same, with the exception that the plates were pre-coated with 0.05 mg/mL poly-D-lysine instead of HFN for MC38 and RAW264.7 cells.

**Immunofluorescence staining and fluorescence microscopy**

After labeling, cells were fixed and permeabilized with cold methanol (pre-chilled at −20 °C) for 10 min at 4 °C. For visualization of BP labeling, cells were washed with PBST (PBS with 0.05% Tween-20) once and then blocked with 1% (w/v) BSA in PBST for 60 minutes at room temperature. To visualize PA labeling, a copper-catalyzed azide–alkyne cycloaddition reaction was performed prior to BSA block and staining. Briefly, cells were incubated with a reaction mixture containing 83.4 μL DPBS, 4 μL Cu/BTTAA complex (12.5 mM $CuSO_4$ and 25 mM BTTAA), 0.6 μL azide-biotin (20 mM stock), and 12 μL freshly prepared sodium ascorbate (5 mg/mL) for 2 h at room temperature. Cells were then washed three times with PBST (Phosphate-Buffered Saline with 0.05% Tween 20) and blocked in 1% BSA (w/v) in PBST for 60 minutes at room temperature. Cells were then incubated with primary antibodies (Mouse anti-V5 antibody, 1:500 dilution, Mouse anti-FLAG antibody, 1:500 dilution; Rabbit anti-TOM20 antibody, 1:400 dilution; Rabbit anti-Calnexin antibody, 1:1000 dilution) in 1% BSA (w/v) in PBST for 2 hours at room temperature. See "Antibodies used in this study" table below for sources. After washing three times with PBST, cells were incubated with secondary antibodies (AlexaFluor488, 1:1000 dilution; AlexaFluor568, 1:1000 dilution; neutravidin-AlexaFluor647, 1:2000 dilution, DAPI, 1:2000 dilution) in 1% BSA (w/v) in PBST for 1 hour at room temperature. Cells were then washed with PBST three times and imaged with a Zeiss AxioObserver inverted microscope with 40x or 63x oil-immersion objective, outfitted with a Yokogawa spinning disk confocal head, a Quad-band notch dichroic mirror (405/488/568/647), and 405 (diode), 491 (DPSS), 561 (DPSS) and 640 nm (diode) lasers (all 50 mW). All images were collected with SlideBook (Intelligent Imaging Innovations) and processed with FIJI.

**RNA extraction and RT-qPCR following biotin-phenol labeling**

After APEX labeling and quenching, cells were gently scraped off the plate and transferred to Eppendorf tubes. Cells were pelleted at 500G for 4 min at room temperature. RNA was extracted from cells using the RNeasy plus mini kit (Qiagen, Cat# 74134) following the manufacturer's protocol. RNA concentrations were determined using Nanodrop.

Enrichment of biotinylated RNA from biotin-phenol labeled samples for RT-qPCR were achieved by following exactly the protocol described in Fazal et al[3]. Briefly, streptavidin magnetic beads (Thermo Fisher Scientific, Cat# 88816) were used at a ratio of 1 μL beads per 2.5 μg of RNA. The beads were washed three times with B&W buffer (5 mM Tris-HCl, pH 7.5, 0.5 mM EDTA, 1 M NaCl, 0.1% TWEEN-20), twice with Solution A (0.1 M NaOH and 0.05 M NaCl), and once with Solution B (0.1 M NaCl). The washed beads were then incubated with RNA samples at 4 °C for 2 hours with rotation.

After incubation, the beads were washed three times with B&W buffer and resuspended in 54 µL of nuclease-free water. Biotinylated RNAs were eluted using a proteinase K-based protocol. A 3× proteinase digestion buffer was prepared containing 330 µL 10× PBS, 330 µL 20% N-Lauryl sarcosine sodium solution, 66 µL 0.5 M EDTA, 16.5 µL 1 M dithiothreitol, and 357.5 µL nuclease-free water. For each reaction, 33 µL of this buffer was added to the beads along with 10 µL of proteinase K (20 mg/mL, Ambion, Cat# AM2548) and 3 µL of Ribolock RNase inhibitor (Thermo Fisher Scientific, Cat# EO0382). The mixture was incubated at 42 °C for 1 hour, followed by 55 °C for 1 hour with shaking. The RNA was subsequently purified using the RNA Clean & Concentrator-5 Kit (Zymo Research, Cat# R1016) according to the manufacturer's instructions. The purified RNA was then prepared for downstream RT-qPCR.

**RNA extraction and RT-qPCR or bulk RNA-seq following phenol azide labeling**

Following phenol-azide labeling and quenching, cells were gently scraped off the plate and pelleted at 500G for 4 min. RNA was extracted using the RNeasy Plus Mini Kit (Qiagen, Cat# 74134). For each condition, a single well from a 6-well plate typically yields 10-30 µg of total RNA and is sufficient for downstream applications. To enrich RNA labeled with phenol-azide for RT-qPCR, 5 µg of total RNA was adjusted to 18.5 µL of 0.5× DPBS (half-strength dilution of 1× Dulbecco's Phosphate-Buffered Saline (DPBS)) was combined with 0.5 µL Ribolock RNase Inhibitor and 1 µL of 2.5 mM DBCO-biotin (Vector Laboratories, Cat# CCT-A105-5, stored in DMSO in -80°C). The reaction was incubated at 37 °C for 1 hour. Following the reaction, excess DBCO-biotin was removed using the RNA Clean & Concentrator-5 Kit (Zymo Research, Cat# R1016). Next, 2.5 µg of total RNA (16 µl) from each group was combined with 4 µl of Superscript IV VILO Master Mix (Thermo Scientific, Cat# 11756050) and subjected to the following thermocycling protocol: 25 °C for 10 min and 50 °C for 10 min. Following reverse transcription, 1 µL of RNA-cDNA hybrids were saved as a pre-pull-down input sample for qPCR analysis. The remaining 19 µL of the RNA-cDNA hybrids were enriched using streptavidin magnetic beads (Thermo Fisher Scientific, Cat# 88816). 1 µL beads were washed twice with B&W buffer (5 mM Tris-HCl, pH 7.5, 0.5 mM EDTA, 1 M NaCl, 0.1% TWEEN-20). The washed beads were resuspended in 20 µL B&W buffer (with 1 µL Ribolock RNase inhibitor (Thermo Fisher Scientific, Cat# EO0382)) and combined with 19 µL RNA-cDNA hybrid samples at room temperature for 30-40 min with gentle rotation. After incubation, the beads were washed once with 100 µL high-salt B&W buffer (5 mM Tris-HCl, pH 7.5, 0.5 mM EDTA, 1 M NaCl, 0.1% TWEEN-20) and once with 100 µL low-salt B&W buffer (5 mM Tris-HCl, pH 7.5, 0.5 mM EDTA, 0.1 M NaCl, 0.1% TWEEN-20). Following the final wash, RNA-cDNA hybrids were directly resuspended in nuclease free $H_2O$ as post-pull-down sample. The on-bead RNA–cDNA complexes were then used directly as templates for qPCR analysis. Transcript recovery was determined as the ratio of RNA amount post-pull-down to the pre-pull-down input using qPCR. qPCR was conducted in 384-well plates using the CFX Connect Real-Time System (Bio-Rad), with a total reaction volume of 10 µl per well. Each reaction consisted of 2.5 µl of diluted cDNA template (pre-pull-down DNA was diluted 1:100, post-pull-down on-bead DNA resuspended in nuclease free $H_2O$ were directly used without further dilution), 2.5 µl of 1 µM forward and reverse primers, and 5 µl of 2× Maxima SYBR Green/ROX qPCR Master Mix (Thermo Scientific, Cat# K0221). The thermal cycling protocol included an initial denaturation step at 95 °C for 3 min, followed by 40 cycles of 95 °C for 10 s and 60 °C for 30 s. A melt-curve analysis was performed from 65 °C to 95 °C, with 0.5 °C increments.

To enrich RNA labeled with phenol-azide for RNA sequencing, 4 µg of total RNA in 18.5 µL of 0.5× DPBS was combined with 0.5 µL Ribolock RNase Inhibitor and 1 µL of 2.5 mM DBCO-biotin (Vector Laboratories, Cat# CCT-A105-5). The reaction was incubated at 37 °C for 1 hour. Following the reaction, excess DBCO-biotin was removed using the RNA Clean & Concentrator-5 Kit (Zymo

Research, Cat# R1016). The entire Clicked RNA sample (~ 4 µg) was used for subsequent streptavidin pull down. Streptavidin magnetic beads (Thermo Fisher Scientific, Cat# 88816) were used at a ratio of 1 µL beads per 5 µg of RNA. The beads were washed twice with B&W buffer (5 mM Tris-HCl, pH 7.5, 0.5 mM EDTA, 1 M NaCl, 0.1% TWEEN-20). The washed beads were then incubated with RNA samples in 40 µL B&W buffer (with 1 µL Ribolock RNase inhibitor (Thermo Fisher Scientific, Cat# EO0382)) at room temperature for 30-40 min with gentle rotation. After incubation, the beads were washed once with 100 µL high-salt B&W buffer (5 mM Tris-HCl, pH 7.5, 0.5 mM EDTA, 1 M NaCl, 0.1% TWEEN-20) and once with 100 µL low-salt B&W buffer (5 mM Tris-HCl, pH 7.5, 0.5 mM EDTA, 0.1 M NaCl, 0.1% TWEEN-20).

Following the final wash, the supernatant was removed, and RNA bound to the beads was fragmented directly in 19.5 µL of Illumina Fragment, Prime, Finish (FPF) buffer (provided in TruSeq Stranded mRNA Library Prep Kit, Illumina, Cat# 20020594) at 94°C for 8 minutes. Immediately after fragmentation (without a 4 °C hold), 17 µL of the supernatant containing the fragmented mRNA was carefully collected for downstream library preparation. Libraries were prepared using the TruSeq Stranded mRNA Library Prep Kit (Illumina, Cat# 20020594). Steps prior to mRNA fragmentation in the library preparation protocol were omitted. From that point forward, the standard manufacturer's protocol was followed, beginning with the combination of 8 µL of the FSA and SuperScript mix with the 17 µL fragmented mRNA. Three modifications were made to the standard procedure: 1) in the "first strand cDNA synthesis" step, Superscript IV Reverse Transcriptase (Thermo Fisher Scientific, Cat# 18090010) was used instead of Superscript II provided in the kit. The corresponding thermal cycler program was modified to: 23°C for 10 minutes, 55°C for 10 min, 80°C for 10 min, hold at 4°C; 2) during the "add index adapter" step, 0.5 µL RNA adapters were used instead of the recommended 2.5 µL; 3) in the "amplify DNA fragment" step, 17 PCR cycles were used for $H_2O_2$ labeled samples, while 20 PCR cycles were used for unlabeled control samples. All remaining steps were performed according to the manufacturer's instructions without modification. In this study, only the TruSeq kit was used for bulk APEX-seq library preparation, and residual rRNA reads were observed. Alternative library preparation kits such as the SMARTer® Stranded Total RNA-Seq Kit v2 – Pico Input Mammalian (Takara, Cat# 634411) incorporates rRNA depletion and is fully compatible with the APEX-seq2 protocol. This workflow begins with RNA fragmentation followed by first strand synthesis, and subsequently removes ribosomal cDNA through ZapR enzyme-mediated cleavage, thereby minimizing rRNA-derived reads.

**Optimization of macrophage polarization conditions**

RAW264.7 cells were polarized into pro-inflammatory macrophages by stimulation with lipopolysaccharide (LPS) (Invivogen, Cat# tlrl-ppglps) and interferon-gamma (IFNγ) (PeproTech, Cat# 315-05), or into anti-inflammatory macrophages using interleukin-4 (IL-4) (PeproTech, Cat# 214-14) and interleukin-13 (IL-13) (PeproTech, Cat# 210-13). Different concentrations of these cytokines were tested to optimize polarization conditions. Following stimulation, the culture media was aspirated, and the cells were washed immediately with cold D-PBS. The cells were then pelleted by centrifugation, and total RNA was extracted using the RNeasy Plus Mini Kit (Qiagen, Cat# 74134). For cDNA synthesis, 1 µg of total RNA (diluted in 8 µL) was combined with 2 µL of Superscript IV VILO Master Mix (Thermo Scientific, Cat# 11756050) and subjected to the following thermal cycling conditions: 25 °C for 10 minutes, 50 °C for 10 minutes, and 85 °C for 5 minutes. The resulting cDNA was then diluted tenfold in nuclease-free water before quantitative PCR (qPCR) analysis. qPCR was performed in 384-well plates using the previously described qPCR protocol. CD86 and Nos2 were selected as pro-inflammatory markers, while CD206 and Arg1 were used as anti-inflammatory markers. Primer sequences are provided in Supplementary Table S3.

**scAPEX-seq and scRNA-seq of co-cultured tumor and macrophage cells**

Raw264.7 cells expressing ERM-APEX2 were seeded at a density of $3 \times 10^5$ cells per well in a 6-well plate. After 24 hours, the culture medium was aspirated, and cells were stimulated with either 10 ng/mL LPS and IFNγ for 8 hours (M1 polarization) or 50 ng/mL IL-4 and IL-13 for 8 hours (M2 polarization). $3 \times 10^5$ MC38 cancer cells expressing ERM-APEX2 were added to stimulated RAW264.7 cells and cocultured for an additional 16 hours. Both cocultured and non-cocultured (M1/M2 state) RAW264.7 and MC38 cells were incubated with 200 μM PA for 30 minutes, followed by the addition of $H_2O_2$ to a final concentration of 1 mM for exactly 1 minute to initiate APEX labeling. The reaction was quenched by adding a 2× quench solution (10 mM Trolox (Sigma Aldrich, Cat# 238813-1G), 20 mM sodium ascorbate (Sigma Aldrich Cat# A7631-25G) and 20 mM sodium azide (Sigma Aldrich, Cat# A7631-25G) in DPBS). Cells were washed three times with DPBS, gently scraped off the plate, and centrifuged at 400 × g for 3 minutes at 4°C. The pellet was resuspended in PBS containing 0.04% (w/v) BSA for single-cell APEX-seq (scAPEX-seq). As a control, conventional single-cell RNA sequencing (scRNA-seq) was performed under identical stimulation and culture conditions, using the same APEX2-ERM-expressing but unlabeled cells. Both cocultured and non-cocultured cells were collected for scRNA-seq.

Single-cell barcoding and GEM (Gel Beads-in-Emulsion) generation were conducted using the 3' v4 kit (10x Genomics, Cat# 1000690), following the manufacturer's protocol with slight modifications: in Step 1.1 (Master Mix Preparation), a custom template switch oligo (TSO) was used instead of the proprietary 10x Genomics TSO, as its biotinylation status is undisclosed. The TSO sequences used were either 5'-/5Me-isodC//iisodG//iMe-isodC/AAGCAGTGGTATCAACGCAGAGTACATrGrGrG-3' (recommended) or 5'-A/isp18/AAGCAGTGGTATCAACGCAGAGTACATrGrGrG-3' (a more cost-effective alternative).[4,5] In both designs, 5'-end modifications were introduced to prevent concatemer formation. The custom TSOs were synthesized by IDT, resuspended in low TE buffer (IDT, Cat# 11-01-02-05) to a final concentration of 1 mM; in Step 1.2, a total of 36,000 cells were loaded; in Step 1.5: the 85°C incubation for 5 minutes during reverse transcription was omitted to preserve RNA-cDNA hybrids; in Step 2.2: 14 PCR cycles used for both the supernatant and the enriched fraction.

Following the addition of the recovery reagent in Step 2.1 (post GEM-RT cleanup), approximately 75 μL of the aqueous phase containing RNA-cDNA hybrids was recovered. To this solution, the following reagents were sequentially added: 7.5 μL of 5 M ammonium acetate (Thermo Fisher Scientific, Cat# AM9070G), 2 μL of Glycogen Blue (15 mg/mL, Thermo Fisher Scientific, Cat# AM9516), 150 μL of cold ethanol. The mixture containing RNA-cDNA hybrids, cold ethanol, glycogen blue and ammomium acetate was incubated at -80°C for 30 minutes, followed by centrifugation at 21,000 × g for 8 minutes at 4°C. The pellet was washed once with 70% ethanol and centrifuged again at 21,000 × g for 5 minutes at 4°C. Residual ethanol was carefully removed, and the pellet was air-dried for 2 minutes before resuspension in 37 μL of buffer EB supplemented with 50 mM NaCl (gentle repeated pipetting is needed to ensure complete dissolution of the pellet). For copper-free click chemistry, the following were added to the resuspended pellet: 1 μL of Ribolock RNase inhibitor, 2 μL of 2.5 mM DBCO-Biotin (Vector Laboratories, Cat# CCT-A105-5). The reaction was incubated at 37°C for 1 hour. To remove excess DBCO-Biotin, the following were added to the solution: 0.5 μL of Glycogen Blue, 4 μL of 5 M ammonium acetate, 120 μL of cold ethanol. The solution was incubated at -80°C for 30 minutes, followed by centrifugation at 21,000 × g for 8 minutes at 4°C. The pellet was washed once with 70% ethanol and centrifuged again at 21,000 × g for 5 minutes at 4°C.

The pellet was resuspended by gentle repeated pipetting until fully dissolved in 35 μL of Elution Solution I (Buffer EB supplemented with 1% Reducing Agent B (10x Genomics, Cat# 2000087) and 0.1% Tween-20, supplemented with 50 mM NaCl and 1 μL of Ribolock RNase inhibitor. Next, 1 µL of pre-washed streptavidin beads, equilibrated in Buffer EB with 50 mM NaCl, was added to capture biotinylated RNA-cDNA hybrids. The mixture was incubated at room temperature with rotation for 40 minutes. After incubation, 35 µL of the supernatant was collected and saved for library preparation. The beads were sequentially washed with 100 µL of high-salt buffer (Buffer EB + 1 M NaCl) and then 100 µL of low-salt buffer (Buffer EB + 50 mM NaCl). Following the washes, the beads were resuspended in 35 µL of Elution Solution I, and the enriched fraction was directly subjected to PCR amplification on the beads, following the protocol outlined in Step 2.2 (cDNA Amplification). All subsequent library preparation steps were performed according to the standard 10x Genomics protocol.

For scRNA-seq sample, single-cell barcoding, GEM generation and library preparation were conducted using the 3' v4 kit (10x Genomics, Cat# 1000690 following the manufacturer's protocol without modification.

Schematic of the experimental setup:

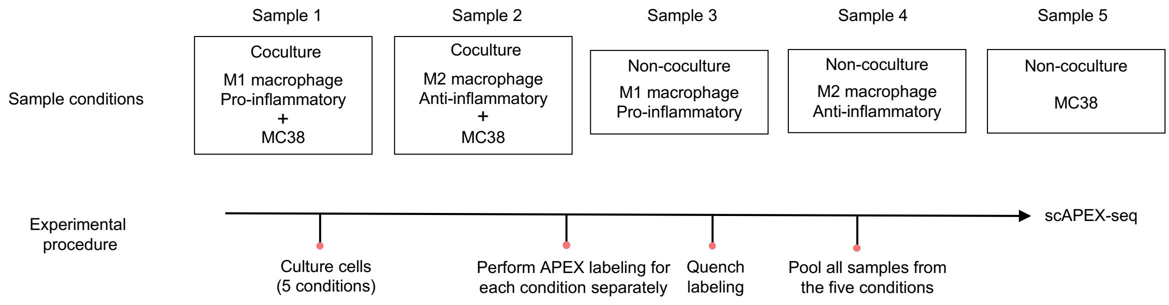


**scAPEX-seq of primary T cells and cancer cells**

Following transduction of HCC1569 cancer cells with ERM-APEX2 and primary CD8+ T cells with both ERM-APEX2 and anti-HER2 scFv, T cells co-expressing the CAR and APEX2 were sorted and expanded for 9 days. During the resting phase, cells were maintained at a density of 0.5 × $10^6$ cells/mL. One day prior to APEX labeling, approximately 1.2 million HCC1569 breast cancer cells expressing ERM-APEX2 were seeded in a 6-well plate. The next day, 0.5 million anti-HER2 scFv $CD8^+$ CAR-T cells expressing ERM-APEX2 and 0.2 million non-CAR $CD8^+$ T cells expressing ERM-APEX2 were added to the cancer cells, centrifuged at 400 × g for 1 minute, and incubated at 37°C in the presence of 100 µM 6-thioguanosine for 1.5 hours. Following this incubation, PA probe was added to a final concentration of 100 µM, and the cells were centrifuged again at 400 × g for 1 minute, followed by an additional incubation at 37°C for 30 minutes. APEX labeling was initiated by adding $H_2O_2$ to a final concentration of 1 mM and incubating for exactly 1 minute. The reaction was immediately quenched by adding 2× quench solution (10 mM Trolox (Sigma Aldrich, Cat# 238813-1G), 20 mM sodium ascorbate (Sigma Aldrich Cat# A7631-25G) and 20 mM sodium azide (Sigma Aldrich, Cat# A7631-25G) in DPBS). T cells from the supernatant and adherent cancer cells were collected by gentle scraping, followed by centrifugation at 300 × g for 3 minutes. The cell pellet was resuspended in quench solution, centrifuged again at 300 × g for 3 minutes, and the supernatant was carefully aspirated. The cells were then resuspended in 1.2 mL of quench solution, transferred to a 1.5 mL tube, and centrifuged. The pellet was resuspended in 700 µL DPBS supplemented with 400 µg/mL BSA, filtered, counted, and kept on ice. For non-cocultured control conditions, HCC1569 cancer cells, CAR-T cells, and non-CAR T cells were labeled separately before pooling them for independent single-cell barcoding and GEM generation. Single-cell barcoding, GEM generation and library preparation were performed using the same scAPEX-seq protocol as described above.

**Chronic antigen stimulation and cell sorting for scAPEX-seq**

HCC1569 cells were seeded at 1 × $10^6$ cells/well in 6-well plates one day prior to the addition of anti-HER2 CAR-T cells expressing APEX-ERM (2 × $10^5$ cells/well). Every three days, cocultures were harvested, centrifuged, and 2 × $10^5$ T cells were re-seeded with freshly plated HCC1569 cells for repeated antigen stimulation. On the seventh stimulation round, target cells were switched to HCC1569 cells expressing ERM-APEX-GFP. After 24 hours of coculture, PA probe was added to a final concentration of 100 µM and incubated at 37°C for 30 minutes. APEX labeling was then initiated by adding $H_2O_2$ (1 mM final concentration) for exactly 1 minute, followed by immediate quenching with 2× quench buffer (10 mM Trolox (Sigma Aldrich, Cat# 238813-1G), 20 mM sodium ascorbate (Sigma Aldrich Cat# A7631-25G) and 20 mM sodium azide (Sigma Aldrich, Cat# A7631-25G) in DPBS). Cells were collected, washed with PBS, and sorted using a BD Aria Fusion cell sorter (BD Biosciences) to enrich APEX-ERM-GFP–expressing T cells and cancer cells separately (see gating strategy in Figure S5G). Sorted populations were processed using modified 10x Genomics 3' v4 protocol as described above.

**NT5E and CTSW CRISPR knock-out**

CRISPR–Cas9 gene knockout in anti-HER2 CAR-T cells was performed by electroporating pre-assembled ribonucleoprotein (RNP) complexes using the P3 Primary Cell 4D Nucleofector X Kit S (Lonza, #V4XP-3032). Two days after activation, 1 × $10^6$ lentiviral transduced CAR-T cells were collected, CD3/CD28 Dynabeads were removed, and cells were pelleted and resuspended in 18 µL of P3 buffer. NT5E RNPs were formed by incubating 1.5 µL of 40 µM Alt-R® S.p. Cas9 Nuclease V3 (IDT, # 1081059) with 300 pmol of NT5E-targeting sgRNA (IDT, Hs.Cas9.NT5E.1.AB) at a 5:1 molar ratio for 15 minutes at 37°C. CTSW RNP were formed by incubating 1.5 µL of 40 µM Alt-R® S.p. Cas9 Nuclease V3 (IDT, # 1081059) with 0.625ul of 150 µM CTSW-targeting sgRNA(Hs.Cas9.CTSW.1.AA) and 0.625ul of 150 µM CTSW-targeting sgRNA(Hs.Cas9.CTSW.1.AC) ratio for 15 minutes at 37°C. The cell suspension was then mixed with the RNPs and electroporated using the EO-115 program in a 16-well cuvette strip. Electroporated cells were immediately transferred into 80 µL of HTCM supplemented with 500 U/mL IL-2, 5 ng/mL IL-7, and 5 ng/mL IL-15, and incubated at at 37°C and 5% CO2 for 10 minutes before expanding the culture to 2 mL. NT5E knockout efficiency was assessed by flow cytometry using an anti-CD73 antibody (Anti-NT5E-PE-Cyanine7, Thermo Fisher Scientific, Cat# 25-0739-42), and CD73 negative cells were sorted to isolate the knockout population (Figure S7C). CTSW knockout efficiency was using Western blot (Figure S8B). Control CAR-T cells were electroporated with a gRNA targeting the safe-harbor locus AAVS1.

**Western blotting**

To assess NT5E protein expression in tumor and T-cell populations, HCC1569 cancer cells and anti-HER2 scFv CD8+ T cells were cocultured for 4 h at a ratio of 1:2. Following coculture, CD8+ T cells were separated from cancer debris by flow sorting. Cocultured or monocultured control cells were lysed with RIPA lysis buffer (Millipore Cat# 20-188) supplemented with 1× Halt Protease Inhibitor Cocktail (Thermo Fisher Scientific, Cat# 78439) and Phenylmethylsulfonyl fluoride (PMSF, MedChemExpress, Cat# HY-B0496). The concentrations of cell lysates were normalized using a BCA Protein Assay Kit (Pierce, Cat# 23225). Proteins were separated by SDS-PAGE on a 10% polyacrylamide gel and subsequently transferred onto a nitrocellulose membrane. To verify protein loading, the membrane was stained with Ponceau S solution (Thermo Fisher Scientific, Cat# A40000279) for 1 minute, then washed with 1× Tris-buffered saline containing 0.1% Tween-20 (TBST, Teknova, Cat# T1688) to remove the stain. The membrane was blocked in 3% BSA in TBST for 30 minutes at room temperature to reduce non-specific binding. After blocking, the membrane was washed three times with TBST, 5 minutes per wash, before incubation with primary antibodies overnight at 4°C in TBST. The primary antibodies used were rabbit anti-NT5E (1:1000 dilution, Cell Signaling Technology, Cat# 13160T) and mouse anti-β-Tubulin (1:2000 dilution, Cell Signaling Technology, Cat# 86298S). Following primary antibody incubation, the membrane was washed three times with TBST and incubated with secondary antibodies for 1 hour at room temperature in TBST. The secondary antibodies used were anti-Mouse IgG Polyclonal Antibody (IRDye® 680RD), 1:2000 dilution (Licor, Cat# 926-68070) and anti-rabbit HRP, (1:2000 dilution, Cell Signaling Technology, Cat# 7074S). The membrane was then washed three times with TBST and imaged using the Odyssey® CLx gel imager (LI-COR Biosciences). To evaluate CTSW expression during repeated antigen stimulation, anti-HER2 scFv $CD8^+$ T cells were cocultured with HCC1569 cancer cells at an E:T ratio of 1:10. Every 3 days, T cells were collected, lysed, and processed for immunoblotting as described above. Primary antibodies used were mouse anti-CTSW (1: 100 dilution, Santa Cruz Biotechnology, Cat# SC-32799) and rabbit anti-GAPDH (1:2000 dilution, Cell Signaling Technology, Cat# 2118S). The secondary antibodies used were anti-Mouse IgG Polyclonal Antibody (IRDye® 680RD) (1:2000 dilution, Licor, Cat# 926-68070) and anti-rabbit IRDye 800CW (1:2000 dilution, LICOR, Cat# 926-32211).

#### ***In vitro*** **T cell cytotoxicity assay for NT5E-KO cells**

HCC1569 cells expressing H2B-mEmerald were seeded at $2 \times 10^5$ cells/well in 24-well plates one day prior to the addition of high-affinity anti-HER2 CAR-T cells (NT5E-knockout or control) at $2 \times 10^4$ cells/well (E:T = 1:10). Every 3 days, cocultures were harvested, centrifuged, and $2 \times 10^4$ T cells were re-seeded with freshly plated HCC1569 cells for repeated antigen stimulation. In parallel, low-affinity anti-HER2 CAR-T cells (NT5E-knockout or control) were seeded at $2 \times 10^5$ cells/well (E:T = 1:1). Plates were imaged every two hours using the Incucyte live-cell imaging system (10× objective, 37°C, 5% $CO_2$), capturing nine fields per well. Cancer cell numbers were quantified based on total integrated green fluorescence intensity, analyzed using the Incucyte's built-in software. Cancer cell killing curves were visualized by plotting normalized total green fluorescence intensity over time.

#### ***In vitro*** **T cell cytotoxicity assay for CTSW overexpressing cells**

HCC1569 cells expressing H2B-mEmerald were seeded at 15,000 cells/well in 96-well plates one day prior to the addition of anti-HER2 CAR-T cells (CTSW overexpression, CTSW knockout or control) at 1,500 cells/well (E:T = 1:10). For CTSW overexpression, primary human CD3+ T cells were cotransduced with anti-HER2 CAR lentivirus and CTSW-T2A-mCherry lentivirus. Control and CTSW knockout cells were cotransduced with anti-HER2 CAR lentivirus and mCherry only lentivirus to ensure matched mCherry expression. Every 3 days, T cells were re-seeded into freshly plated HCC1569 cells (30,000 cells/well) for repeated antigen stimulation. Plates were imaged every two hours using the Incucyte live-cell imaging system (10× objective, 37°C, 5% $CO_2$), capturing nine fields per well. Cancer cell numbers were quantified based on total integrated green fluorescence intensity, analyzed using the Incucyte's built-in software. T cells numbers were counted by flow cytometry after each round. Cancer cell killing curves were visualized by plotting total green fluorescence intensity over time.

#### **Flow cytometry and cell counting**

All flow cytometry and cell counting were performed using the Novocyte Quanteon Flow Cytometer. CAR expression levels were assessed using anti-Myc antibody (Cell signaling, Cat# 2233S). T cell differentiation status was evaluated using anti-CD45RA (Biolegend, Cat# 304140) and anti-CD62L (BioLegend, Cat# 304822) antibodies. Surface IL-2 receptor alpha (CD25) levels were measured with anti-CD25 antibody (BioLegend, Cat# 356114).

For CAR-T and cancer cell co-culture experiments, following each round of antigen stimulation, cells were first stained with a Live/Dead fixable green viability dye (Thermo Fisher, Cat#

L34969, 1:1000, in PBS) according to the manufacturer's protocol. Subsequently, live T cells were stained with either anti-CD8 (Invitrogen, Cat# 46-0087-42) or anti-CD4 (biolegend, Cat# 980812) antibodies. Cell counts were obtained using the instrument's built-in absolute count function. Unless otherwise specified, all antibodies were diluted in 1:100 in PBS supplemented with 5% FBS.

**Statistical analysis**
Gaphpad Prism 10 (GraphPad Software) was used for the analysis of data and generation of bar graphs. Statistical significance was assessed using unpaired two-tailed t-tests for comparisons involving a single factor or two-way analysis of variance (ANOVA) for experiments involving two factors.

**Bulk APEX-seq data analysis**
Transcript abundances for all samples were quantified from raw reads using *kallisto (v0.50.1)*[6]. Gene level abundances were then calculated by summing over transcript abundances. Low count genes were filtered out by requiring at least one sample to have 10 or more counts. Differential expression analysis was then performed using DESeq2 (PyDESeq2 implementation)[7,8]. Cytosolic and secretory RNA annotations were derived from Villenueva et al.[9], using enrichment value cutoffs of 0.4 and 0.8 respectively. The set of mitochondrial RNAs were defined as the 13 protein coding genes in the mitochondrial genome. Long non-coding RNAs were used as the nuclear-specific RNA set, derived from Werner et al[10] using chromatin association ratio > 1.5. The set of OMM-associated RNAs is derived from the original APEX-seq study[3], using OMM_Log$_2$FC > 1.

**scAPEX-seq data preprocessing**
For all scAPEX-seq libraries, reads were demultiplexed using *bcl2fastq2*. Resulting fastq files were processed using Cell Ranger v8.0.1 (10x Genomics). Cell Ranger filtered feature barcode matrices were used as input for downstream analysis. Downstream analysis was performed in scanpy v1.10.4[11]. Outlier cells in total counts, total genes detected, and percent mitochondrial reads were removed. Genes detected in too few cells were also removed. Doublets were detected and removed using scrublet[12]. Denoised counts were generated using the scAR software package[13]. Total count normalization and log transformation was applied to the counts. Afterwards, standard feature selection and dimensionality reduction methods, as outlined in the scanpy documentation, were performed.

**Single-cell APEX-seq-macrophage-cancer coculture data analysis**
UMAP visualizations were used to assign cluster labels based on well characterized M1, M2, and MC38 cell-type markers[14,15]. For specificity analysis, we again used RNA annotations derived from Villenueva et al.[9], by mapping from human genes to mouse homologs, and calculate scAPEX-seq enrichment as the mean ratio of scAPEX-seq counts to Supernatant-seq counts across the sample for a given cell-type. MT and OMM genes (same gene lists as in bulk APEX-seq) are provided as a negative control for enrichment. For sensitivity analysis, we use the same scAPEX-seq enrichment ratio and gene list for ER membrane associated RNAs, using Log2FC > 0.5 as a cutoff. Cell matching between scAPEX-seq and Supernatant-seq was performed using direct matching of cell barcodes between samples.

Predicted CCIs were generated using the rank_aggregate method LIANA+[16] to aggregate CCI predictions across multiple methods. Rank_aggregate was run using default settings, and filtered using cellphoneDB generated p-values (p-val < 0.05 cutoff). CCIs were filtered for uniqueness (by ligand + receptor complex identity) for comparisons between clusters.

Datasets were integrated using the Scanorama software package[17], also using default parameters. For any downstream analysis that involved comparison of expression between samples, Scanorama batch correction was also applied. For DEG analysis, diffxpy v0.7.4 (https://github.com/theislab/diffxpy) was used. For scRNA-seq DEG analysis, for each cell-type, differences between the two largest clusters were analyzed. For scAPEX-seq, the comparison was performed between the coculture and non-coculture clusters for each cell-type.

#### Single-cell APEX-seq-CAR-T-cancer coculture data analysis

DEG analysis, cell cluster cell-type identification, CCI analysis, and data integration were performed as described above. For 2hr coculture CAR-T experiment, secretory differential enrichment was calculated by calculating DEGs for memory CAR-T cells vs. noncoculture CAR-T cells in both scAPEX-seq and Supernatant-seq, and then taking the difference in Log2FC between the scAPEX-seq and Supernatant-seq per each gene. CellOracle[18] analysis was performed according to standard procedures as outlined in the CellOracle documentation, treating scAPEX-seq and Supernatant-seq as independent scRNA-seq datasets.

We applied *dynamo*[19] for transcriptomic vector analysis in our scAPEX-seq and Supernatant-seq datasets. Spliced and unspliced counts were quantified using velocyto[20]. Scanorama-integrated CAR-T data across samples was used to calculate cell velocities, generate the vector field, and also for reanalyzing vector fields with *in silico* gene perturbation. For secretory velocity, Supernatant-seq data was used as a direct substitute for unspliced reads, while spliced reads were substituted with scAPEX-seq data.

#### Code availability

Codes to generate each figure are available on https://github.com/alicetinglab/scAPEXseq.

#### Data availability

All sequencing data are available through the Gene Expression Omnibus (GEO) under accession GSE315660, GSE315659.

#### Key resources table

| REAGENT or RESOURCE | SOURCE | IDENTIFIER |
|---|---|---|
| Antibodies | | |
| Anti-V5 | Thermo Fisher Scientific | Cat# R96025; RRID: AB_2556564 |
| Anti FLAG | Agilent | Cat# 200472; RRID: AB_10596649 |
| Anti-TOMM20 | Abcam | Cat# Ab186735; RRID: AB_2889972 |
| Anti-Calnexin | Thermo Fisher Scientific | Cat# PA534754; RRID: AB_2552106 |
| Anti-rabbit-AlexaFlour405 | Thermo Fisher Scientific | Cat# A-31556; RRID: AB_221605 |
| Anti-mouse-AlexaFlour488 | Thermo Fisher Scientific | Cat# A-11029; RRID: AB_2534088 |
| Anti-mouse-AlexaFluor568 | Thermo Fisher Scientific | Cat# A-21134; RRID: AB_2535773 |
| NeutravidinAlexaFluor647 | Thermo Fisher Scientific | Cat# A2666 |
| Streptavidin IRDye 800CW | LI-COR | Cat# 926-32230 |
| Anti-Mouse IRDye 680RD | LI-COR | Cat# 926-68070 |
| Anti-Rabbit IRDye 800CW | LI-COR | Cat# 926-32211 |

| | | |
|---|---|---|
| Anti-Rabbit-HRP | Cell Signaling Technology | Cat#7074S |
| Anti-NT5E | Cell Signaling Technology | Cat#13160T |
| Anti-NT5E-PE-Cyanine7 | Thermo Fisher Scientific | Cat# 25-0739-42; RRID: AB_2573368 |
| Anti-β-Tubulin | Cell Signaling Technology | Cat# 86298S |
| Anti-Myc | Cell Signaling Technology | Cat# 2233S |
| DAPI | Enzo Life Sciences | Cat# AP402-0010 |
| Anti-mouse-CTSW | Santa Cruz Biotechnology | Cat# SC-32799 |
| Anti-rabbit-GAPDH | Cell signaling technology | Cat# 2118S |
| Anti-CD45RA | Biolegend | Cat# 304140 |
| Anti-CD62L | Biolegend | Cat# 304822 |
| Anti-CD25 | Biolegend | Cat# 356114 |
| Anti-CD4 | Biolegend | Cat# 980812 |
| Anti-CD8 | Invitrogen | Cat# 46-0087-42 |

Chemicals, peptides and recombinant proteins

| | | |
|---|---|---|
| Sodium ascorbate | Sigma Aldrich | Cat# A7631-25G |
| Trolox | Sigma Aldrich | Cat# 238813-1G |
| Sodium azide | Sigma Aldrich | Cat# S2002-5G |
| Biotin tyramide | APEXBio | Cat# A8011 |
| AMPure XP Beads | Beckman Coulter | Cat# A63881 |
| Pierce streptavidin magnetic beads | Thermo Fischer Scientific | Cat# 88816 |
| Fibronectin | Millipore | Cat# FC010 |
| RiboLock RNase inhibitor | Thermo Fisher Scientific | Cat# EO0384 |
| Puromycin | VWR | Cat# 80054-138 |
| Superscript IV Reverse Transcriptase | Thermo Fisher Scientific | Cat# 18090010 |
| Proteinase K solution (20 mg/ml) | Thermo Fisher Scientific | Cat# AM2548 |
| Hydrogen peroxide solution, 30% (w/w) | Sigma Aldrich | Cat# H1009-100ML |
| DBCO-biotin | Vector Laboratories | Cat# CCT-A105-5 |
| Phenol azide | Sai Life Sciences | N/A |
| LPS | Invivogen | Cat# tlrl-ppglps |
| IFN-γ | PeproTech | Cat# 315-05 |
| IL-4 | PeproTech | Cat# 214-14 |
| IL-13 | PeproTech | Cat# 210-13 |
| Polyethylenimine (PEI) | Polysciences | Cat# 24765 |
| Human anti-CD3/CD28 Dynabeads | Life Technologies | Cat# 11131D |
| RPMI 1640 Medium | Gibco | Cat# 11875135 |
| X-VIVO® 15 Serum-free Hematopoietic Cell Medium | Lonza | Cat# 04-418Q |
| N-acetyl L-cysteine | Sigma-Aldrich | Cat# A9165 |
| CELLBANKER 1 | Zenoaq | Cat# 11910 |
| Trans-LT1 | Mirus Bio | Cat# MIR 2300 |
| Opti-MEM | Thermo Fisher Scientific | Cat# 31985062 |
| Polybrene | Thermo Fisher Scientific | Cat# 725-2545 |

Critical commercial assays

| | | |
|---|---|---|
| Superscript IV Vilo Master Mix | Thermo Fischer Scientific | Cat# 11756050 |
| RNeasy Plus mini kit | QIAGEN | Cat# 74134 |
| RNA Clean and concentrator-5 | Zymo Research | Cat# R1016 |
| TruSeq Stranded mRNA Library Prep Kit | Illumina | Cat# 20020594 |
| Chromium GEM-X Single Cell 3' Kit v4 | 10x Genomics | Cat# PN-1000690 |
| RosetteSep™ Human CD8+ T Cell Enrichment Cocktail | STEMCELL Technologies | Cat# 15063 |

Deposited data

| | | |
|---|---|---|
| Raw and analyzed data | This paper | All sequencing data are available through the Gene Expression Omnibus (GEO) under accession GSE315660, GSE315659 |

Experimental models: Cell lines

| | | |
|---|---|---|
| HEK293T | ATCC | Cat# CRL-3216 |
| HCC1569 | ATCC | Cat# CRL-2330 |
| MC-38 | Kerafast | Cat# ENH204-FP |
| Raw264.7-ASC | Invivogen | Cat# raw-asc |
| Lenti-X 293T cells | Takara | Cat# 632180 |

Recombinant DNA

| | |
|---|---|
| ERM-APEX2 | Addgene Plasmid #79055 |
| APEX2-OMM | Addgene Plasmid #79056 |
| APEX2-NES | Addgene Plasmid #92158 |
| Mito-APEX2 | Addgene Plasmid #72480 |
| APEX2-NLS | Addgene Plasmid #124617 |
| psPAX2 | Addgene Plasmid #12260 |
| pMD2.G | Addgene Plasmid #12259 |
| pPCVd8.91 | Addgene Plasmid #187441 |
| pHR_SFFV | Addgene Plasmid #79121 |
| pHR_SFFv_4D5-High-CAR_RHL002 | Addgene Plasmid #164825 |
